# K-nearest neighbor smoothing for high-throughput single-cell RNA-Seq data

**DOI:** 10.1101/217737

**Authors:** Florian Wagner, Yun Yan, Itai Yanai

## Abstract

High-throughput single-cell RNA-Seq (scRNA-Seq) is a powerful approach for studying heterogeneous tissues and dynamic cellular processes. However, compared to bulk RNA-Seq, single-cell expression profiles are extremely noisy, as they only capture a fraction of the transcripts present in the cell. Here, we propose the k-nearest neighbor smoothing (kNN-smoothing) algorithm, designed to reduce noise by aggregating information from similar cells (neighbors) in a computationally efficient and statistically tractable manner. The algorithm is based on the observation that across protocols, the technical noise exhibited by UMI-filtered scRNA-Seq data closely follows Poisson statistics. Smoothing is performed by first identifying the nearest neighbors of each cell in a step-wise fashion, based on partially smoothed and variance-stabilized expression profiles, and then aggregating their transcript counts. We show that kNN-smoothing greatly improves the detection of clusters of cells and co-expressed genes, and clearly outperforms other smoothing methods on simulated data. To accurately perform smoothing for datasets containing highly similar cell populations, we propose the kNN-smoothing 2 algorithm, in which neighbors are determined after projecting the partially smoothed data onto the first few principal components. We show that unlike its predecessor, kNN-smoothing 2 can accurately distinguish between cells from different T cell subsets, and enables their identification in peripheral blood using unsupervised methods. Our work facilitates the analysis of scRNA-Seq data across a broad range of applications, including the identification of cell populations in heterogeneous tissues and the characterization of dynamic processes such as cellular differentiation. Reference implementations of our algorithms can be found at https://github.com/yanailab/knn-smoothing.

## INTRODUCTION

Over the past decade, single-cell expression profiling by sequencing (scRNA-Seq) technology has advanced rapidly. After the transcriptomic profiling of a single cell (Tang et al. 2009), protocols were developed that incorporated cell-specific barcodes to enable the efficient profiling of tens or hundreds of cells in parallel (Islam, Kjällquist, et al. 2011; Hashimshony, Wagner, et al. 2012). scRNA-Seq methods were then improved by the incorporation of unique molecular identifiers (UMIs) that allow the identification and counting of individual transcripts (e.g., Islam, Zeisel, et al. 2014; Hashimshony, Senderovich, et al. 2016). More recently, single-cell protocols were combined with microfluidic technology (Dijk et al. 2017; Macosko et al. 2015; Zheng et al. 2017), combinatorial barcoding (Cao et al. 2017; Rosenberg et al. 2017), or nanowell plates (Gierahn et al. 2017). These high-throughput scRNA-Seq methods allow the cost-efficient profiling of tens of thousands of cells in a single experiment.

Due to the typically very low amounts of starting material, and the inefficiencies of the various chemical reactions involved in library preparation, scRNA-Seq data is inherently very noisy (Ziegenhain et al. 2017). This has motivated the development of many specialized statistical models, for example for determining differential expression (Kharchenko, Silberstein, and Scadden 2014), performing factor analysis (Pierson and Yau 2015), pathway analysis (Fan et al. 2016), or more general modeling of scRNA-Seq data (Risso et al. 2017). In addition, methods have been proposed to impute missing values (W. V. Li and J. J. Li 2017) and to perform smoothing (Dijk et al. 2017). Finally, many authors of scRNA-Seq studies have relied on ad-hoc approaches for mitigating noise, for example by clustering and averaging cells belonging to each cluster (Shekhar et al. 2016; Baron et al. 2016).

Fundamental to any statistical treatment are the assumptions that are made about the data. For methods aimed at analyzing scRNA-Seq data, assumptions about the noise characteristics determine which approach can be considered the most appropriate. All aforementioned approaches have assumed an overabundance of zero values, compared to what would be expected if the data followed a Poisson or negative binomial distribution. However, in the absence of true expression differences, the analysis by Ziegenhain et al. (2017) has suggested that across scRNA-Seq protocols, there is little evidence of excess-Poisson variability when expression is quantified by counting unique UMI sequences (“UMI filtering”) instead of raw reads (see Figure 5B in Ziegenhain et al. (2017)). This is consistent with reports describing individual UMI-based scRNA-Seq protocols, which have demonstrated that in the absence of true expression differences, the mean-variance relationship of genes or spike-ins closely follows that of Poisson-distributed data (Grün, Kester, and Oudenaarden 2014; Dijk et al. 2017; Zheng et al. 2017).

**Figure 5.**
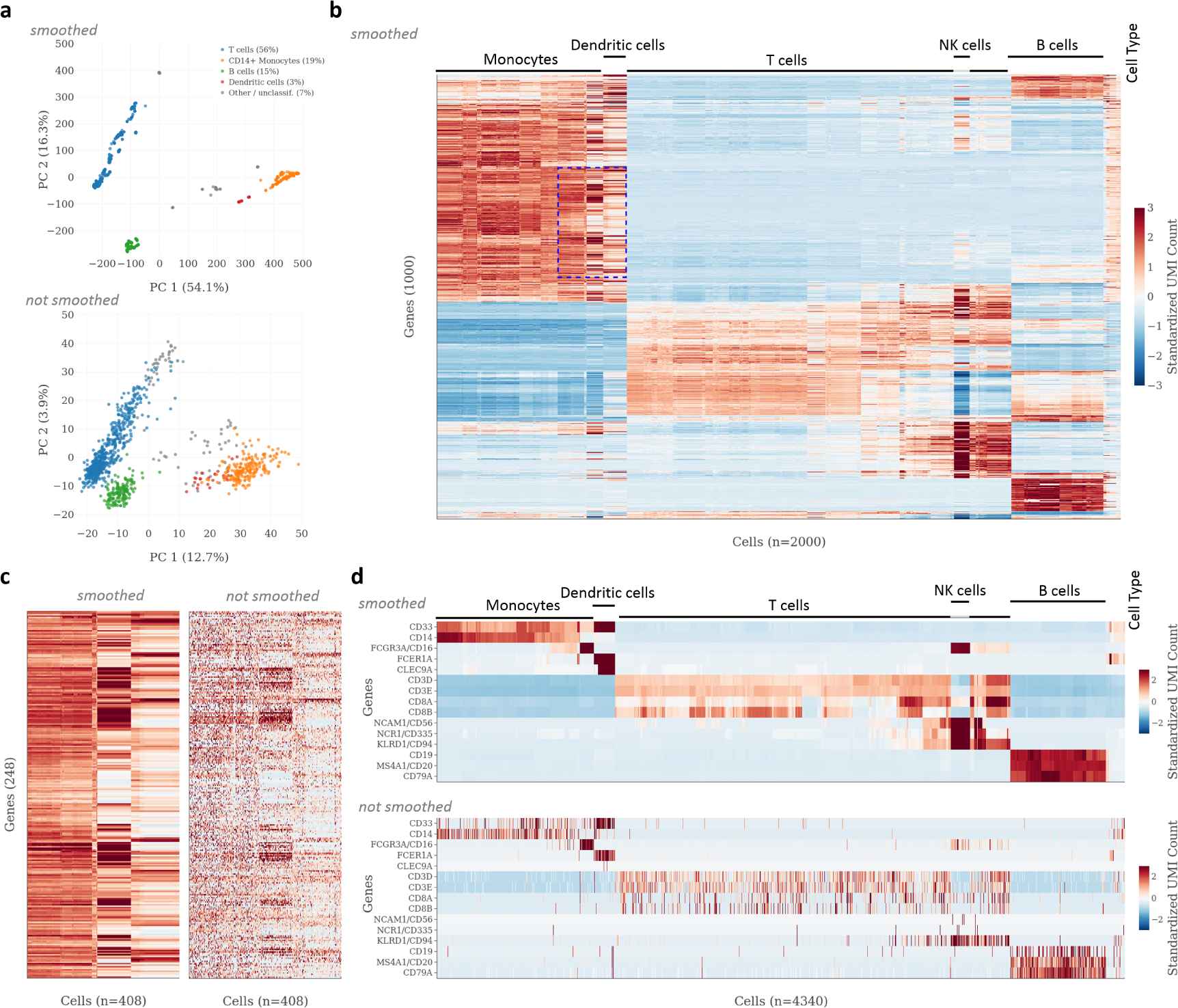
Application of k-nearest neighbor smoothing to scRNA-Seq data from human peripheral blood mononuclear cells (PBMCs) All panels show data from the `PMBC` dataset, published online by 10x Genomics. **a**-**c** Panels showing results of PCA and hierarchical clustering on smoothed and unsmoothed data, as in Figure 4. Cell types in (**a**) were identified based on the smoothed data, using ad-hoc expression thresholds for a list of marker genes compiled from the literature (see Methods). T cells were defined as having expression of *CD83D* ≥ 500 TPM (UMI-filtered transcripts per million); CD14+ monocytes, *CD14* ≥ 250 TPM; B cells, *CD79A* ≥ 1,000 TPM; dendritic cells, *FCER1A* ≥ 500 TPM. Cells that exceeded none of the thresholds, or more than one, were labeled as “other / unclassified”. Due to technical limitations of the visualization library used, only a random subset of 2,000 cells (out of the 4,340 cells in the dataset) is shown in (**b**). **d** Expression of selected marker genes for the major cell types present in the data, with (top) and without (bottom) smoothing.

In this work, we propose two smoothing algorithms that make direct use of the observation that after normalization to account for efficiency noise (Grün, Kester, and Oudenaarden 2014), the technical noise associated with UMI counts from high-throughput scRNA-Seq protocols is entirely consistent with Poisson statistics, implying that the observed transcripts for each cell represent a small, random sample of all the transcripts present. Instead of developing a parametric model, we propose algorithms that smooth scRNA-Seq data by aggregating gene-specific UMI counts from the *k* nearest neighbors of each cell. To accurately determine these neighbors, we propose to proceed in a step-wise fashion using partially smoothed profiles, and to make use of an appropriate variance-stabilizing transformation. Conveniently, the noise associated with the smoothed expression values is again Poisson-distributed, which simplifies their variance-stabilization and downstream analysis. We demonstrate the improved signal-to-noise ratio of scRNA-Seq data processed with our algorithms on real-world examples, and quantitatively compare the accuracies of the smoothing methods proposed here and elsewhere (Dijk et al. 2017; W. V. Li and J. J. Li 2017) on simulated scRNA-Seq data.

## RESULTS

### The normalized UMI counts of replicate scRNA-Seq profiles are Poisson-distributed

To validate the Poisson-distributed nature of high-throughput scRNA-Seq data in the absence of true expression differences, we obtained data from control experiments conducted on three platforms: in-Drop (Dijk et al. 2017), Drop-Seq (Macosko et al. 2015), and 10x Genomics (Zheng et al. 2017). In these experiments, droplets containing identical RNA pools were analyzed. Assuming that the number of transcripts in each droplet was sufficiently large, there are no true expression differences among droplets, and all of the observed differences among droplets can be attributed to technical noise arising from library preparation and sequencing. As expected from published results (cf. Figure 5A in Klein et al. (2015), Supplementary Figure 2f in Zheng et al. (2017)), data from both the inDrop platform and the 10x Genomics platform followed the Poisson distribution (see Figure 1a,c; see Methods), with the exception of highly expressed genes, which is likely due to global droplet-to-droplet differences in capture efficiency, previously referred to as “efficiency noise” (Grün, Kester, and Oudenaarden 2014).

**Figure 1.**
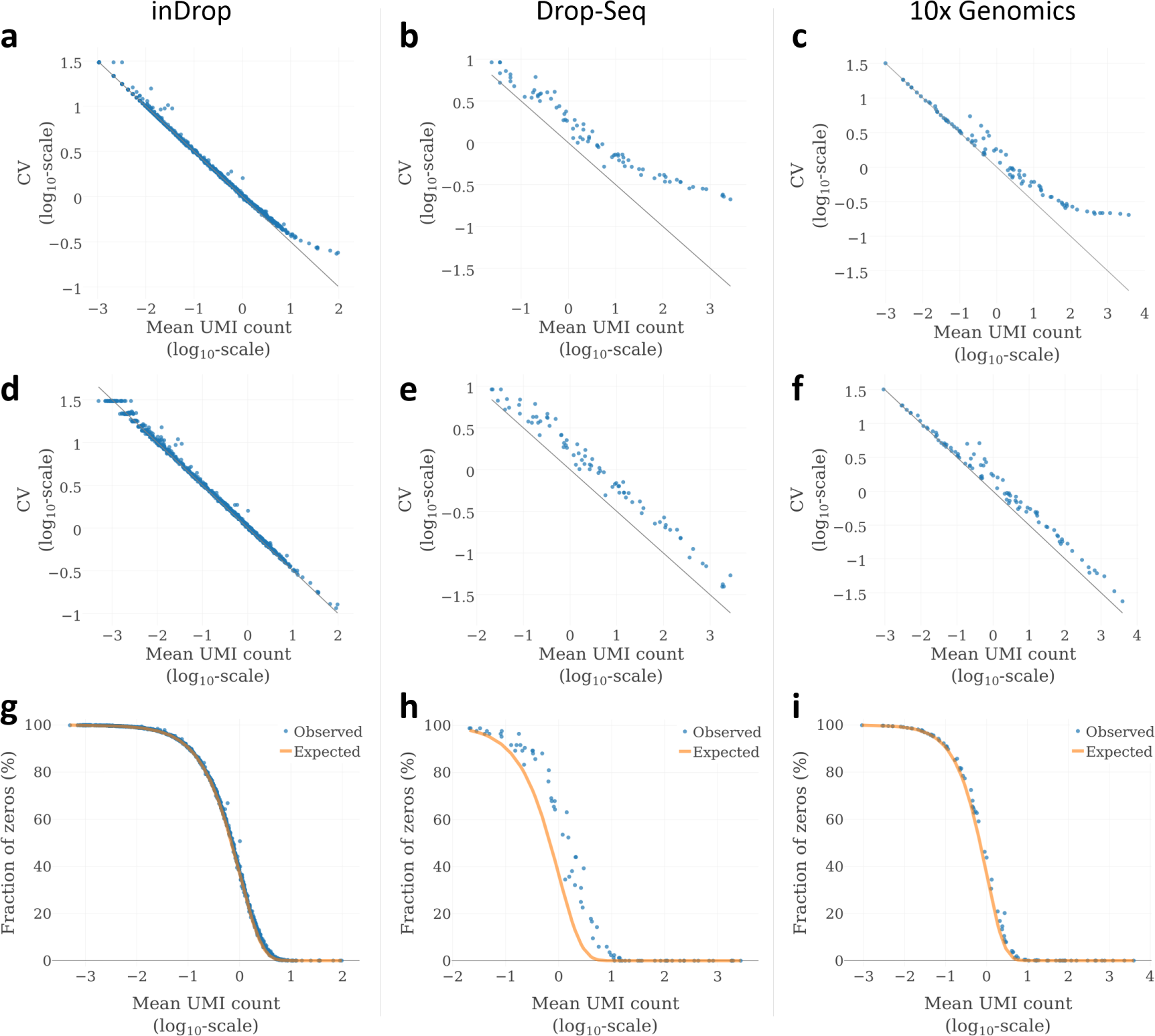
Noise profiles of three high-throughput single-cell RNA-Seq platforms. (**a-c**) Relationship between mean UMI count and coefficient of variation (CV) in pure RNA replicates, analyzed using inDrop (**a**) Drop-seq (**b**), and 10x Genomics (**c**). For inDrop, RNA was extracted from cultered cells (Dijk et al. 2017). For Drop-Seq and 10x Genomics, ERCC spike-in RNA was analyzed (see Macosko et al. (2015) and Zheng et al. (2017)). (**d-f**) The same relationship after normalizing each profile to the median total UMI count (see Methods). (**g-i**) Expected vs. observed fraction of zeros, as a function of mean expression (after median-normalization). For inDrop data (**a**, **d** and **g**), a randomly sampled subset of 1,000 genes is shown for better readability.

**Figure 2.**
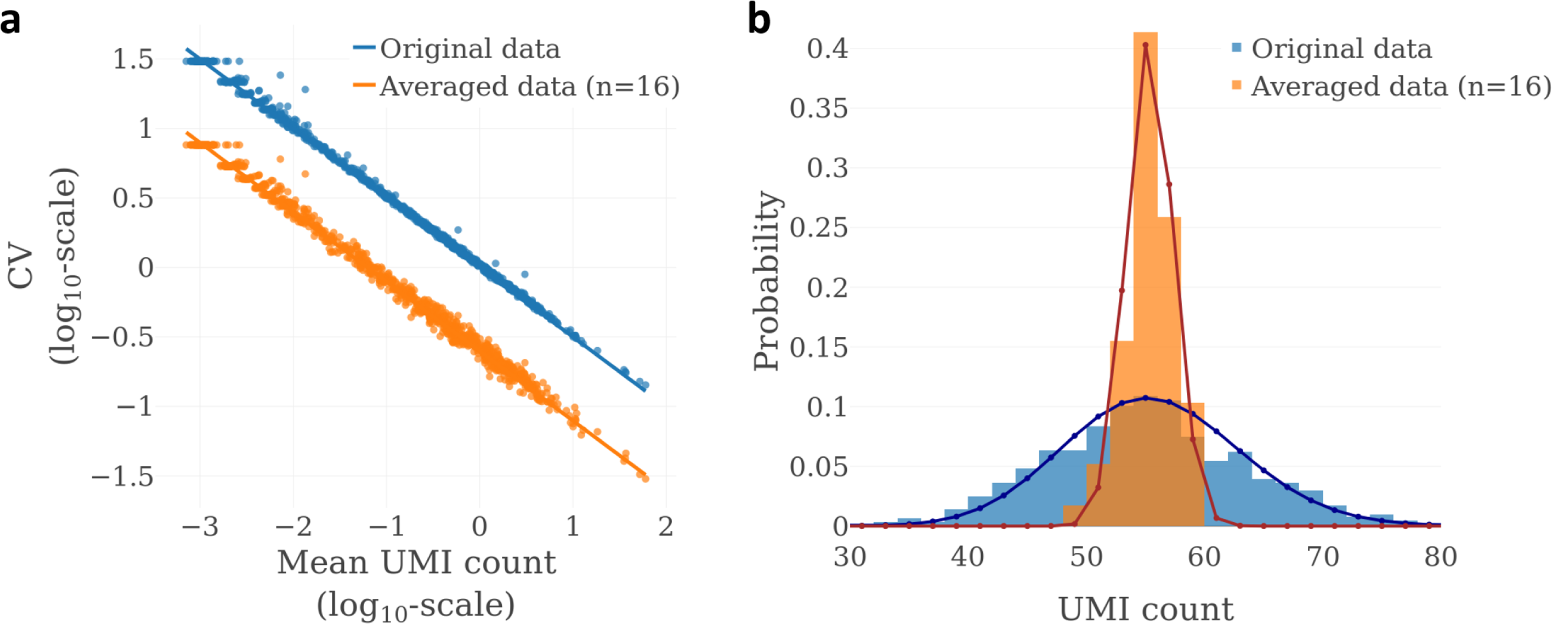
Simple averaging of scRNA-Seq expression profile replicates reduces the coefficient of variation in a manner predicted by Poisson statistics. (**a**) Effect of averaging on the coefficient of variation, for 1,000 randomly selected genes in the inDrop pure RNA dataset (Klein et al., 2015). Solid lines represent the theoretical relationship based on the Poisson distribution. After averaging of 16 profiles at a time, the CV can be seen shifted downwards by about 0.6 units, which corresponds to a factor of 4 on the log_10_-scale used. (**b**) Distribution of UMI counts for the *GAPDH* gene, before and after averaging. Bars show the observed UMI distributions. The solid lines show the theoretical distributions for a Poisson-distributed variable representing the original values (blue), and a scaled Poisson-distributed variable representing the averaged values (orange). To eliminate efficiency noise, both original and averaged profiles were normalized to the median total UMI count (Grün et al., 2014).

For the Drop-Seq data, Macosko et al. (2015) did not discuss the mean-variance relationship, but we observed a pattern consistent with inDrop and 10x Genomics data (see Figure 3b). Interestingly, the y axis intercept of the Drop-Seq CV-mean relationship was clearly above 0, suggesting that transcript counts followed a scaled Poisson distribution (see Methods). A possible explanation could be that the computational pipeline used to derive the Drop-Seq UMI counts generated artificially inflated transcript counts, but we did not explore this hypothesis further.

**Figure 3.**
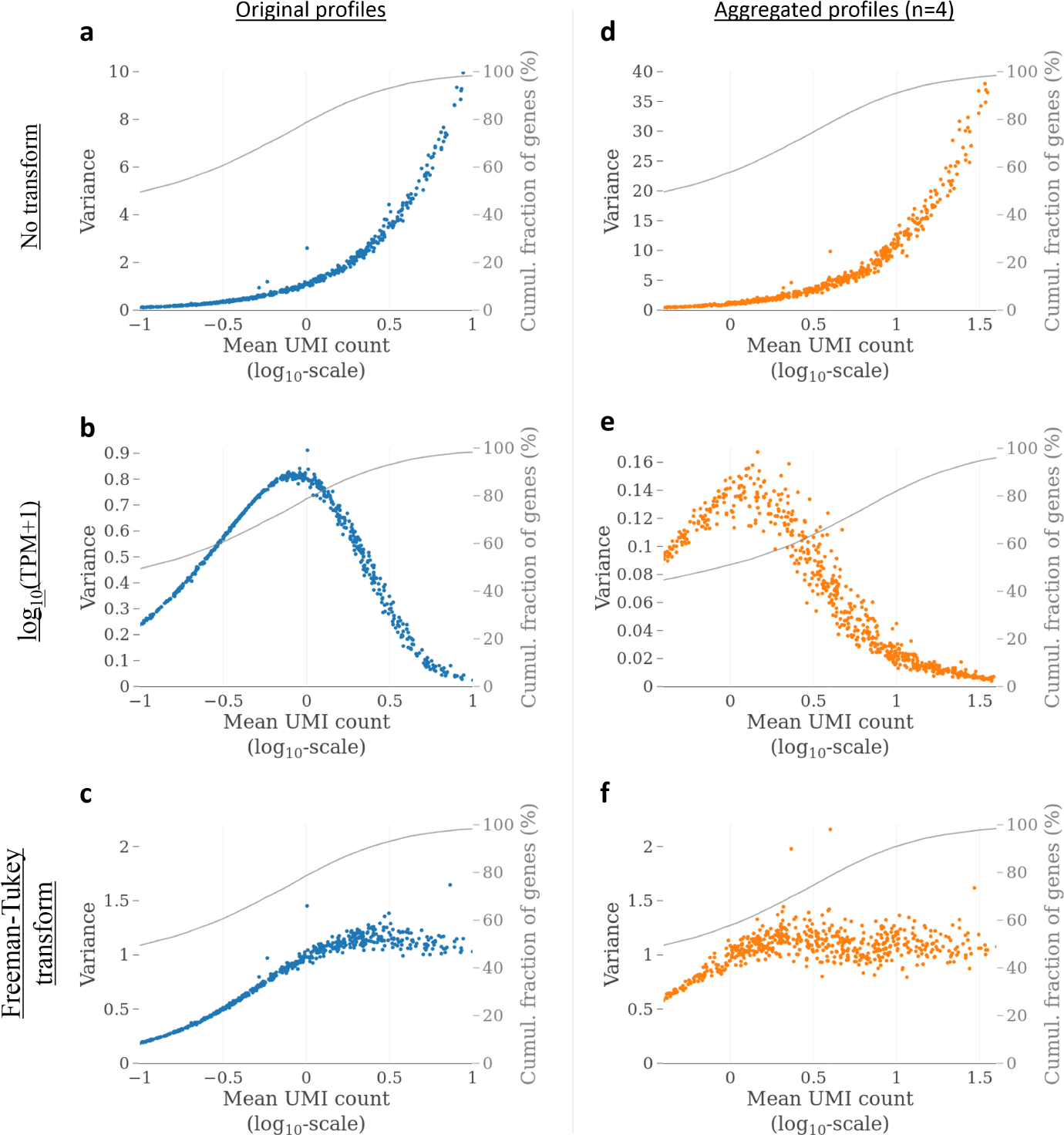
Effect of scRNA-Seq data transformations on mean-variance relationships in technical replicates from the inDrop protocol. All data are normalized to the median total UMI count. (**a-c**) Gene mean-variance relationships in the pure RNA samples (Klein et al., 2015) without transformation, with log_10_(TPM+1) transform, and with Freeman-Tukey transform 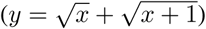, respectively. (**d-f**) Mean-variance relationships after aggregating the expression profiles of randomly selected, non-overlapping batches of 4 cells, for the same transformations. All plots show data for the same 1,000 randomly selected genes.

To test whether the larger-than-expected variance of highly expressed genes can indeed be explained by efficiency noise, we normalized the expression profiles in each dataset to the median UMI count across profiles (Model I in Grün, Kester, and Oudenaarden (2014); see Methods). This resulted in an almost perfectly linear CV-mean relationship (see Figure 1d-f), suggesting that efficiency noise is indeed the dominating source of variation for very highly expressed genes.

Finally, we directly compared the frequency of UMI counts of zero for each gene to that predicted by Poisson statistics, and found that for the inDrop and 10x Genomics data, the observed values matched the theoretical prediction almost perfectly (see Figure 3g,i). For the Drop-Seq data, the frequency of zeros was slightly shifted upwards across the entire expression range (see Figure 3h), which may be due to artificially inflated UMI counts (see Methods).

In summary, we found that for all three high-throughput scRNA-Seq platforms examined, Poisson-distributed noise, in combination with the efficiency noise observed for very highly expressed genes, described virtually all of the observed technical variance, and that there was no evidence of substantial zero-inflation. We note that the recent publication describing the Quartz-Seq2 single-cell platform also reports a Poisson noise relationship (see Figure 2e in Sasagawa et al. (2017)), bringing the total number of high-throughput scRNA-Seq protocols with reported Poisson noise characteristics to four.

### Aggregation of *n* replicate profiles results in Poisson-distributed values with the signal-to-noise ratio increased by a factor of 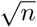

Since the sum of independent Poisson-distributed variables is again Poisson-distributed, we reasoned that the aggregation of normalized expression values from *n* independent measurements of the same RNA pool would result in Poisson-distributed values, with the signal-to-noise ratio increased by a factor of 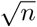 (see Methods). Similarly, we predicted that averaging instead of aggregating (summing) would result in a scaled Poisson distribution with the same increased signal-to-noise ratio. We tested this idea on the inDrop pure RNA dataset previously shown in Figure 1a, which consisted of 935 expression profiles. Averaging randomly selected, non-overlapping sets of 16 profiles resulted in 58 new expression profiles, with genes exhibiting an almost exact four-fold increase in their signal-to-noise ratios, i.e., a four-fold reduction of their coefficients of variation, as expected (see Figure 2a). As an example, the UMI count distribution of the *GADPH* gene before and after averaging is shown in Figure 2b, and can be seen to closely match the theoretically predicted Poisson and scaled Poisson distributions, respectively. In summary, the results showed that independently of gene expression level, aggregating expression values from replicate profiles led to more accurate expression estimates that again exhibited Poisson-distributed noise profiles.

### The Freeman-Tukey transform effectively stabilizes the technical variance of high-throughput scRNA-Seq data

Based on the aforementioned results, we conceived an algorithm to smooth single-cell RNA-seq data, with the following outline:

- For each cell *C*:
  1. Determine the *k* nearest neighbors of *C*.
  2. Calculate a smoothed expression profile for *C* by combining its UMI counts with those of the *k* nearest neighbors, on a gene-by-gene basis.
  3. (Optional) Divide *C*’s new expression profile by *k* + 1, to retain the scale of the original data.

The main challenge in implementing this algorithm is to devise an appropriate approach for determining the *k* nearest neighbors of each cell, and to choose an appropriate *k*. We defer the question of how to choose *k* to the Discussion, and focus here on the problem of determining the *k* nearest neighbors.

Due to the Poisson-distributed nature of scRNA-Seq data, the technical variance (noise) associated with each gene is directly proportional to its expression level. This type of extreme heteroskedasticity poses a problem when attempting to calculate cell-cell similarities, because the noise of highly expressed genes can drown out the true expression differences of more lowly expressed genes, therefore strongly biasing the analysis towards the most highly expressed genes. One strategy to address this issue is the application of an appropriate variance-stabilizing transformation, designed to render the technical variance independent of the gene expression level (Love, Huber, and Anders 2014). For bulk RNA-Seq data, a log-TPM (or log-RPKM) transform is commonly used for this purpose, even though lowly expressed genes will still exhibit unduly large variances under this transformation (Love, Huber, and Anders 2014). Based on our results, we reasoned that for scRNA-Seq data, the *Freeman-Tukey transform* (FTT), 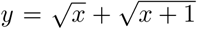, would be a more appropriate choice, as it is designed to stabilize the variance of Poisson-distributed variables (Freeman and Tukey 1950).

To compare the abilities of the FTT and the log-TPM (transcripts per million) transform to stabilize the technical variance of scRNA-Seq data, we applied both transformations to the inDrop pure RNA dataset, and found that the FTT produced significantly better results (see Figure 3): With the log transform, genes with low-intermediate expression, which we considered to be those with expression values between the 60th and 80th percentile rank (of all protein-coding genes, not only genes expressed by K562 cells), had between three- and ten-fold higher levels of variance than the 10% most highly expressed genes (see Figure 3b). In contrast, with the FTT, the difference was no larger than two-fold, and the variances of lowly expressed genes were biased downwards, not upwards (see Figure 3c). Moreover, we found that the FTT also stabilized the variance of the aggregated profiles (see Figure 3d-f), which was expected, given our earlier observation that the aggregated UMI counts are again Poisson-distributed. In particular, a greater share of genes now had variances close to 1. This closely mirrored theoretical results, according to which the variance Poisson-distributed variables with mean *λ* ≥ 1 should be within 6% of the asymptotic value of 1 after FTT (Freeman and Tukey 1950). In summary, our analysis showed that distance calculations performed on Freeman-Tukey transformed (FT-transformed) UMI counts would give similar weight to genes with intermediate and high expression. Expression differences from lowly expressed genes would tend to be suppressed, but this suppression would become less severe for aggregated expression profiles.

### A k-nearest neighbor smoothing algorithm for scRNA-Seq data

The previously discussed ideas suggested that a simple way to determine the *k* nearest neighbors for all cells would be to normalize their expression profiles, apply the FTT, and then find the *k* closest cells for each cell using the Euclidean distance metric. However, we reasoned that this simple approach could be improved upon, because the noisiness of the data itself can interfere with the accurate determination of the *k* nearest neighbors. We therefore instead decided to adopt a step-wise approach, whereby initially, each profile is only minimally smoothed (using *k*_1_ = 1). In the second step, a larger set of nearest neighbors (e.g., *k*_2_ = 3) is identified for each cell based on those minimally smoothed profiles, and the raw data is then smoothed using these larger sets of neighbors. Additional steps using increasing *k*_*i*_ are performed until the desired degree of smoothing is reached (i.e., *k*_*i*_ = *k*). By choosing the *i*’th step to use *k*_*i*_ = min{2^*i*^ − 1, *k*}, each step theoretically improves the signal-to-noise ratio of each individual expression measurement by a factor of 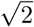 — except for the last step, for which the improvement can be smaller —, and only a small number of steps are required even for large choices of *k* (e.g., six steps for *k* = 63). The resulting “kNN-smoothing” algorithm is formalized in Algorithm 1 (see https://github.com/yanailab/knn-smoothing for reference implementations in Python, R, and Matlab). Using simulation studies, we found that in contrast to a simple “one-step” algorithm, the step-wise approach resulted in a significantly more accurate selection of neighbors, especially for large *k* (see below).

### Application of kNN-smoothing to scRNA-Seq data of human pancreatic islets improves clustering results and recovers specific expression patterns for marker genes

To test whether kNN-smoothing would improve the ability to distinguish between different cell types in a scRNA-Seq experiment, we applied the algorithm (with *k*=15) to a single-cell expression dataset obtained from human pancreatic islet tissue, containing at least 14 distinct cell populations (Baron et al. 2016) (`PANCREAS` dataset). We first performed principal component analyses (PCA; see Methods) and observed several improvements after smoothing (see Figure 4a): First, cell type clusters appeared significantly more compact in principal component space, indicating that the smoothed expression profiles were more similar than unsmoothed profiles for cells of the same type, but more different for cells from distinct types. Second, a single cluster of cells that contained alpha cells as well as other cells separated into two highly distinct clusters after smoothing. Notably, all alpha cells were still contained within a single cluster after smoothing. This suggested smoothing helped reveal important differences that were not previously captured by the first two principal components. Third, the proportion of cells of each type that could be identified using simple marker gene expression thresholds increased slightly, suggesting that the expression values of individual marker was less noisy in the smoothed data. Finally, a much greater share of total variance was explained by the first two principal components (PCs) for the smoothed data than for the unsmoothed data (40.3% vs 20.8%), which is consistent with a greater share of variance originating from true biological differences rather than technical noise.

**Figure 4.**
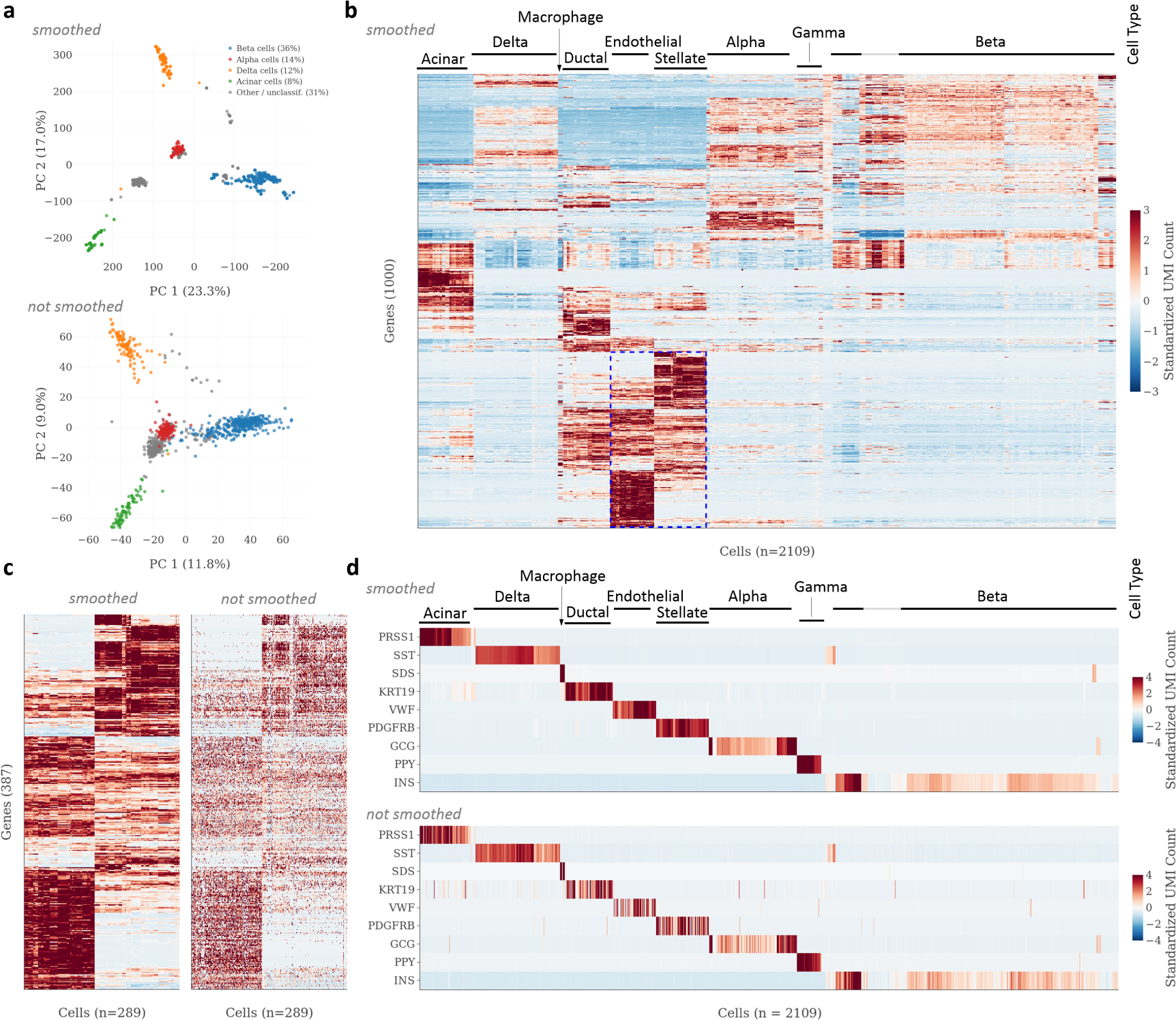
Application of k-nearest neighbor smoothing to scRNA-Seq data from human pancreatic islet tissue. All panels show data from the `PANCREAS` dataset, from a study by Baron et al. (2016). Smoothing was performed using *k* = 15. **a** Principal component analysis (PCA) with (top) and without (bottom) smoothing. Axis labels indicate the fraction of variance explained. Cell types were identified based on the smoothed data, using ad-hoc expression thresholds for the marker genes listed in Baron et al. (2016). Beta cells were defined as having expression of *INS* ≥ 40,000 TPM (UMI-filtered transcripts per million); alpha cells, *GCG* ≥ 5,000 TPM; delta cells, *SST* ≥ 20,000 TPM; acinar cells, *CPA1* ≥ 1,000 TPM. Cells that exceeded none of the thresholds, or more than one, were labeled as “other / unclassified”. **b** Heatmap showing clustered and standardized expression data for the 1,000 most variable genes, after smoothing. **c** Heatmap providing a zoomed-in view of the area marked in blue in (**b**), with (left) and without (right) smoothing. **d** Expression of cell type-specific marker genes (Baron et al. 2016) with (top) and without (bottom) smoothing. Cells are ordered as in (**b**). See Methods for details on how PCA and hierarchical clustering were performed.

We next performed hierarchical clustering on the smoothed data after filtering for the 1,000 most variable genes (see Methods). When we visualized the results as an expression heatmap (Eisen et al. 1998), several gene and cell clusters were readily discernible (see Figure 4b). A direct comparison between the smoothed and unsmoothed data showed that smoothing produced significantly less noisy expression patterns while preserving expression differences between relatively similar cell populations (see Figure 4c). To assess whether cell clusters delineated different cell types, we examined the expression patterns of known marker genes for nine cell types present in the data (Baron et al. 2016), and found that the hierarchical clustering of the smoothed expression profiles accurately grouped cells by their cell type (see Figure 4d, top panel). Moreover, compared to the unsmoothed data, the expression patterns of these marker genes appeared significantly less noisy (see Figure 4d, bottom panel). Finally, we repeated the entire analysis on the unsmoothed data, and found that it was considerably more difficult to discern clusters of genes and cells (see Figure S1a), and that judging by the expression patterns of the marker genes, not all cell types were clustered together appropriately (see Figure S1b). In summary, our analyses showed that kNN-smoothing with *k*=15 significantly improved the results obtained with PCA as well as hierarchical clustering, and that it recovered stable and cell type-specific expression patterns for all of the marker genes examined.

#### Algorithm 1 K-nearest neighbor smoothing for UMI-filtered scRNA-Seq data

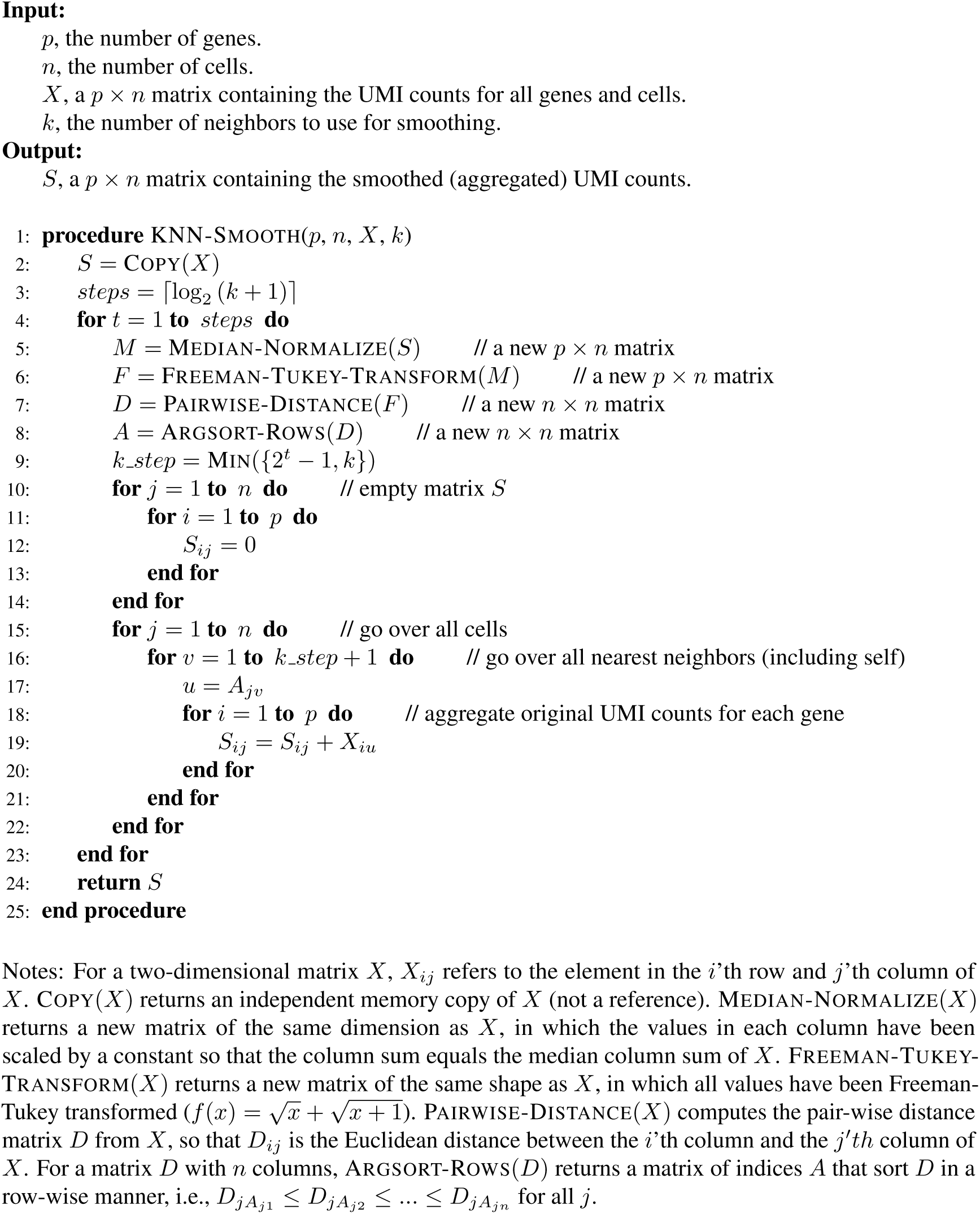

### Application of kNN-smoothing to scRNA-Seq data of human peripheral blood mononuclear cells recovers robust expression profiles for diverse immune cell populations

As a second test of our algorithm, we applied kNN-smoothing to a dataset containing scRNA-Seq data for 4,340 peripheral blood mononuclear cells (PBMCs), obtained using the 10x Genomics “Chromium” protocol (the `PMBC` dataset; see Methods). PBMCs can easily be obtained from peripheral blood, have been studied extensively, and contain a diverse set of immune cell types (Kleiveland 2015), thus enjoying popularity as a point of reference for scRNA-Seq studies (e.g., Zheng et al. 2017; Gierahn et al. 2017). The identification and characterization of immune cell types in peripheral blood using scRNA-Seq is also an activate area of investigation (e.g., Villani et al. 2017). Since the `PMBC` dataset contained significantly more cells than the `PANCREAS` dataset, and the expression profiles exhibited significantly higher complexity (i.e., expression levels were less concentrated on a few highly expressed genes; data not shown), we chose to apply more aggressive smoothing using *k*=127. We compared the results of PCA applied before and after smoothing, and found that, again, smoothing significantly improved the compactness of cell type clusters in principal component space, and strongly increased the fraction of variance explained by the first two PCs — this time, from 16.6% to 70.4%. Moreover, using expression thresholds for individual marker genes (see below), we were able to assign one of four major cell type identities (T cells, CD14 monocytes, B cells, and dendritic cells) to 93% of all cells in the smoothed data. The unidentified cells likely included NK cells as well as technical outliers.

Next, we performed hierarchical clustering after filtering for the 1,000 most variable genes, visualized the results as a heatmap, and obtained several easily distinguishable clusters of cells and genes, providing an overview of the heterogeneity in the data (see Figure 5b). Repeating the same clustering procedure on the unsmoothed data produced much less coherent clusters (see Figure S2). We compared the smoothed and smoothed data within a small region of the heatmap in a side-by-side comparison and observed that smoothing dramatically reduced the apparent noise levels, while largely preserving differences between similar sets of cells (see Figure 5c). Finally, we compiled a list of marker genes for the major cell types found in PBMC samples, including T cells, monocytes, B cells, NK cells, and dendritic cells (see Methods). In comparing the expression patterns of these genes across cells ordered according to the hierarchical clustering results, we found that smoothed resulted in vastly more stable expression patterns, while the expression of each marker gene remained confined to a specific subset of cells. A comparison with the full heatmap suggested that within most cell types, there existed significant population substructure. For example, several distinct clusters of cells were apparent among the set of T cells expressing *CD3D* and *CD3E*, which likely distinguish specific subsets such as CD4 and CD8 T cells, or naive and memory T cells. In summary, the application of aggressive smoothing (with *k*=127) to PBMC data led to significant improvements in the ability to cluster cells by their cell type, and produced stable and cell type-specific specific expression patterns for marker genes, thus demonstrating the applicability of kNN-smoothing to data generated using 10x Genomics’ high-throughput scRNA-Seq solution.

### Comparison with other smoothing methods on simulated datasets shows superior performance of kNN-smoothing

To quantitatively compare the accuracy of kNN-smoothing with that of other smoothing methods, we devised an approach for simulating scRNA-Seq datasets containing a mixture of cell types. Our idea was to base each simulation on a real scRNA-Seq dataset, in order to make the simulated data as similar to real scRNA-Seq expression data as possible, both biologically and technically. To ensure biological similarity, we simulated clusters with expression profiles obtained from the real data, based on hierarchical clustering results. To ensure technical fidelity, we simulated Poisson-distributed sampling noise, modeled on top of efficiency noise, the distribution of which was again obtained from the real data (see Methods for details). We generated two datasets, `SIM-PANCREAS` (based on the `PANCREAS` dataset) and `SIM-PBMC` (based on the `PMBC` dataset). A visual comparison based on clustered heatmaps illustrated the similarity between real and simulated scRNA-Seq data (see Figures S3 and S4). We then applied kNN-smoothing, MAGIC (Dijk et al. 2017), and scImpute (W. V. Li and J. J. Li 2017) to the two datasets, and quantified the similarity of the results to the true cluster profiles from which the cell expression profiles were generated.

We tested different parameter settings for each method, and observed that as expected, the choice of *k* had a large effect on the accuracy of the results obtained with kNN-smoothing (see Figure 6). However, for all values of *k* ≥ 15 that we tested (up to *k*=511), kNN-smoothing outperformed MAGIC and scImpute on both datasets by a large margin, independently of the way in which we quantified accuracy. We first quantified the relative accuracy of each cell’s expression profile by calculating its Pearson correlation coefficient (PCC) with the true cluster expression profile, on log_2_-transformed data. For kNN-smoothing with *k*=15, the median PCC across all cells in the `SIM-PANCREAS` dataset was approx. 0.93. For *k*=63, it was approx. 0.98. In contrast, the best values obtained by MAGIC and scImpute across all parameter settings were approx. 0.85 and 0.87, respectively (see Figure 6a). These differences were even more pronounced for the SIM-PBMC dataset (see Figure 6c), and when we quantified absolute accuracies by root-mean squared error (RMSE) on log-transformed data (see Figure 6b,d). We then quantified accuracies, using both PCC and RMSE, on square root-transformed data instead of log_2_-transformed data. This resulted in slightly smaller absolute differences, but we again observed that kNN-smoothing clearly outperformed the other methods for *k* ≥ 15 (see Figure S5).

**Figure 6.**
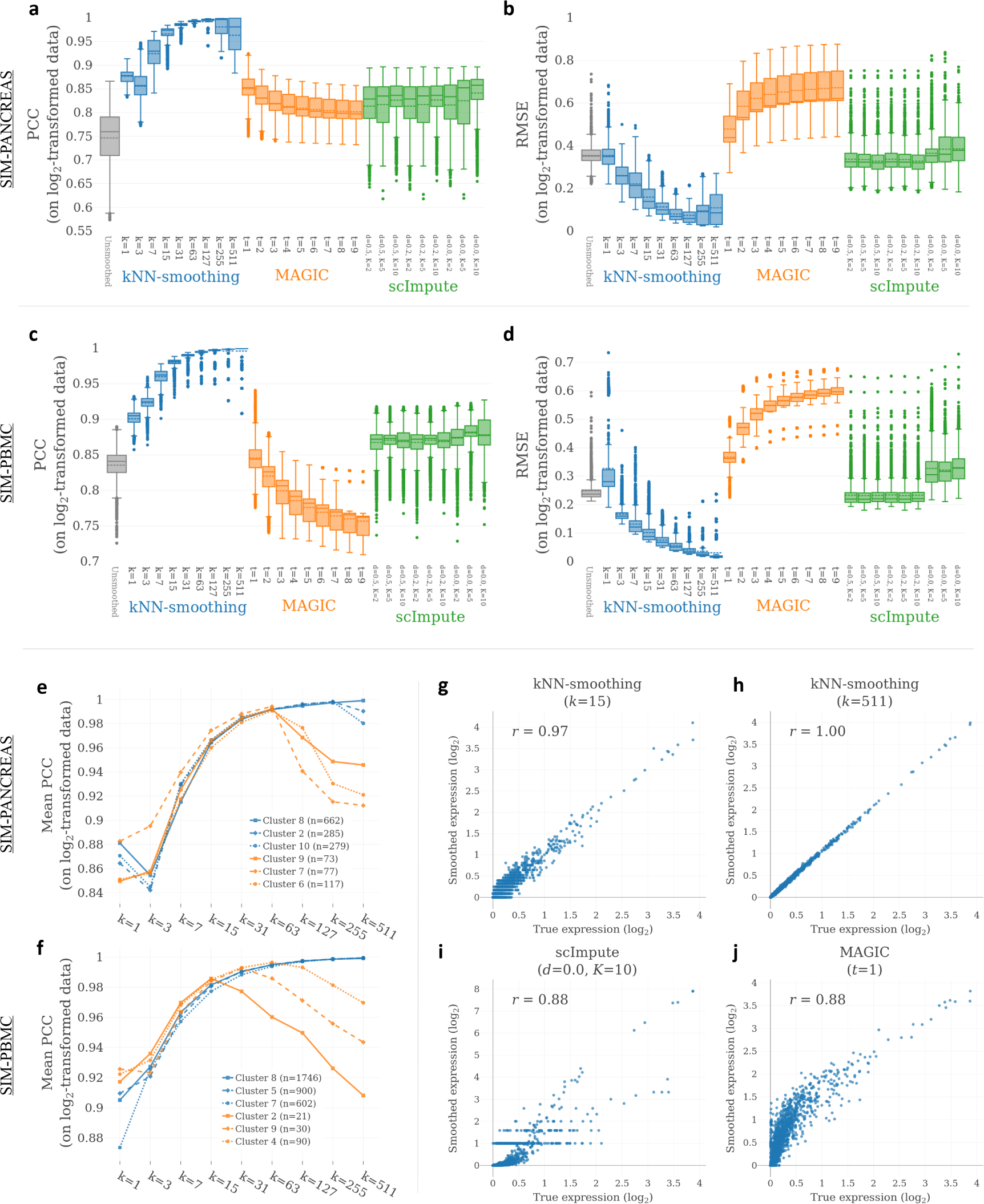
Accuracy of kNN-smoothing in comparison to other smoothing methods for simulated scRNA-Seq data. Accuracy on `SIM-PANCREAS` dataset. **c, d**. Accuracy on `SIM-PBMC` dataset. (**a**) and (**c**) show relative accuracy of log_2_-transformed expression profiles, quantified using the Pearson correlation coefficient (PCC). (**b**) and (**d**) show absolute accuracy of log_2_-transformed expression profiles, quantified using root mean squared error (RMSE). Box plots summarize the distributions of values for all cells in the data. The three methods were each run with various different parameter settings, indicated on the x-axis (see Methods for details). **e,f** Average accuracy (PCC) of cells in the three largest and smallest clusters of the `SIM-PANCREAS` dataset (**e**) and `SIM-PBMC` (**f**) dataset, respectively, for different settings of *k* as indicated on the x-axis. **g-j** Correlation between true and smoothed expression profile for a representative cell from the largest cluster in the `SIM-PANCREAS` dataset, for kNN-smoothing, scImpute, and MAGIC, with parameter settings indicated above each panel.

Our evaluation of kNN-smoothing on simulated data also showed that up to a certain point, choosing larger values of *k* produced increasingly accurate expression profiles. In fact, the median PCC for *k*=511 was very close to 1 in the SIM-PBMC dataset (see Figure 6c). However, the best median PCC for the `SIM-PANCREAS` dataset was obtained for *k*=255, and a significant fraction of cells exhibited much lower accuracies for *k*=255 and *k*=511 compared to *k*=127 (see Figure 6a). This apparent “over-smoothing” was not surprising, since a significant fraction of cells in the `SIM-PANCREAS` dataset belonged to clusters that were represented by less than 256 cells. Therefore, some of the 255 neighbors selected for these cells had to belong to other clusters, and using their expression values for smoothing resulted in less accurate expression profiles. To confirm that cluster size determined whether or not cells benefitted from smoothing with very large *k*, we examined the average accuracies of cells from the three largest and smallest clusters for different *k*. In both datasets, we observed that as predicted, accuracies started to drop off whenever *k* was chosen larger than the cluster size (see Figure 6e,f).

To obtain a more detailed view of the results of kNN-smoothing, MAGIC, and scImpute, we selected a representative cell from the largest cluster in the `PANCREAS` dataset (*n*=662), and examined the correlation of the smoothed profiles with the true cluster profile using scatter plots. For kNN-smoothing, we examined the results for *k*=15 and *k*=511, whereas for MAGIC and scImpute, we picked the parameter settings that achieved the best median PCC across all cells. The correlations for this particular cell mirrored the overall results (see Figure 6g-j), which showed that kNN-smoothing with either setting of *k* produced more highly correlated profiles than either of the two other methods. However, whereas the PCC for both MAGIC and scImpute was 0.88, the values reported by MAGIC were merely noisy and non-linear, while the scImpute results also exhibited some obvious smoothing artifacts (see Figure 6j).

Finally, we observed that for *k*=3, the median PCC of kNN-smoothing was sometimes lower than that for *k*=1. We believe this surprising result is related to size biases by the algorithm in the selection of neighbors (cells) to be used for smoothing (further discussed below). In conclusion, our evaluation of different smoothing methods on two simulated datasets showed that kNN-smoothing outperformed the other methods by a large margin for most choices of *k*, and in some cases recovered cell expression profiles with near-perfect accuracy.

### Other variants of kNN-smoothing are less accurate and exhibit stronger size selection bias in simulated datasets

In the design of our smoothing algorithm, we made several decisions based on theoretical considerations, as well as our intuitions. We therefore aimed to examine whether the performance of the resulting algorithm retrospectively validated these decisions. Specifically, we aimed to compare the kNN-smoothing algorithm to a variant in which neighbors are identified in a single step, as opposed to a step-wise approach. Second, we aimed to test whether the choice of calculating cell-cell distances on median-normalized and FT-transformed data performed better than using the more commonly employed TPM normalization, followed by a log-transformation. We refer two these two variants as the “single-step” variant and the “log-TPM” variant, respectively.

To test the accuracy of the different variants of the smoothing algorithm, we again relied on our simulated datasets (see above), and determined, for a range of different *k*, the fraction of cells with incorrect neighbors for each variant. We found that the log-TPM variant performed very poorly in both datasets, resulting in approximately 80% and 20%, respectively, of cells having an incorrect neighbor even for *k* = 1 in `SIM-PANCREAS` and SIM-PBMC (see Figure 7a,b). The “one-step” variant performed generally worse than the step-wise variant, with the exception of *k* = 15 and *k* = 31 in the `SIM-PBMC` dataset.

**Figure 7.**
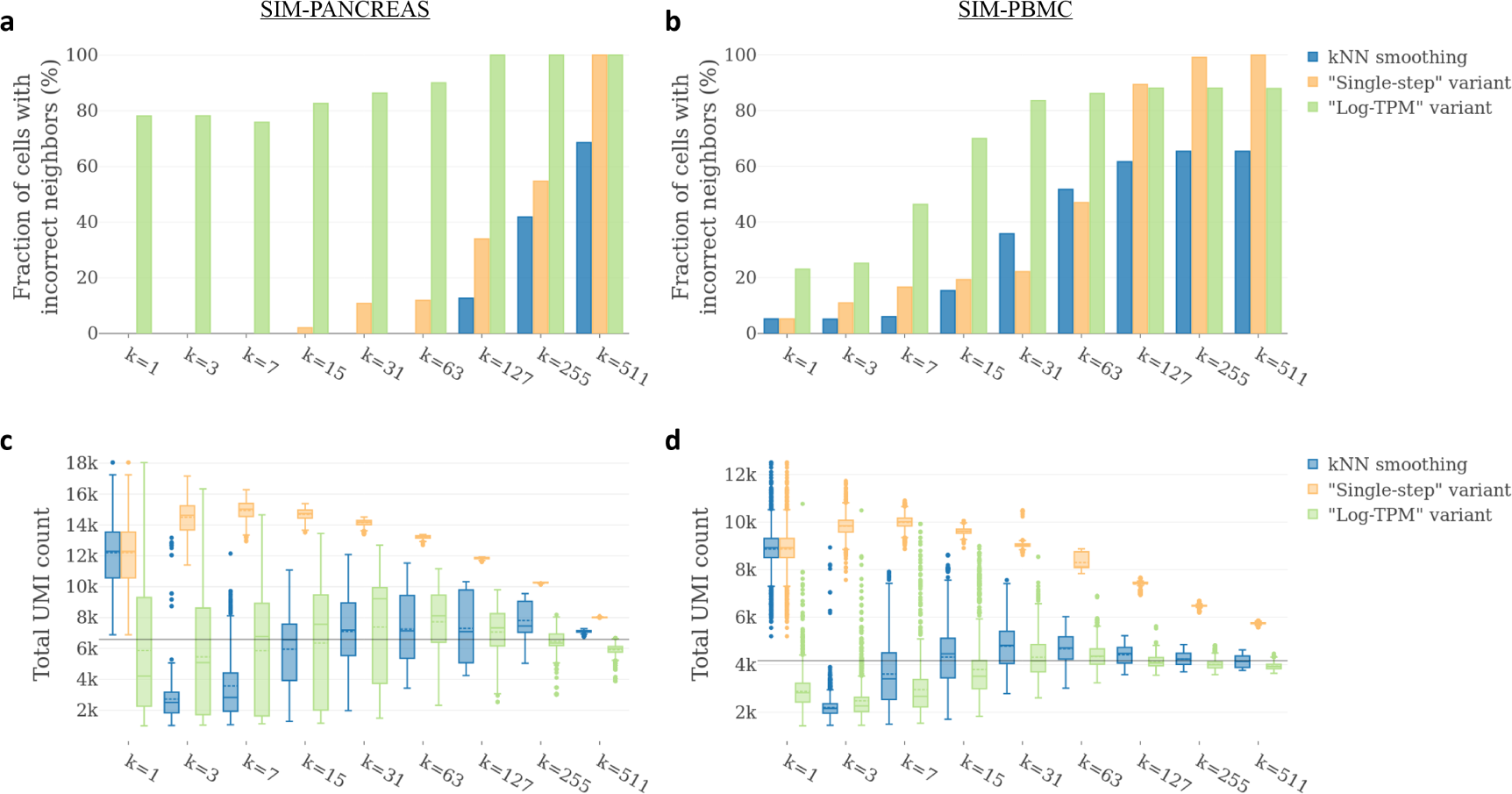
Accuracy and size bias of kNN-smoothing in comparison to two variants of the algorithm, for simulated scRNA-Seq data. **a, b** Accuracy quantified as the fraction of cells with “incorrect” neighbors selected by the smoothing algorithm when applied to the `SIM-PANCREAS` (**a**) and `SIM-PBMC` (**b**) datasets, respectively, with different settings of *k*, as indicated on the x-axis. A cell has an “incorrect neighbor” when at least one cell “neighbor” from a different cluster was included in the calculation of its smoothed expression profile. **c, d** Size bias measured by the total UMI count per cell in the `SIM-PANCREAS` (**c**) and `SIM-PBMC` (**d**) datasets, respectively, after smoothing with different settings of *k*, as indicated on the x-axis.

Over the course of our simulation experiments, we noticed that the average “sizes” (total UMI counts) of the smoothed “cells” (expression profiles) sometimes deviated significantly from the true UMI count of each cluster, which could only be explained by a size bias in the way in which neighbors were selected for each cell (the sizes of cells belonging to the same cluster varied due to our simulation of efficiency noise; see Methods). To examine whether kNN-smoothing and the two variants exhibited different size biases, we compared the distribution of smoothed profile sizes for a range of different *k*, focusing only on cells from the largest cluster in each dataset (see Figure 7c,d). We found that the algorithms exhibited strikingly different behaviors. Most notably, the one-step variant exhibited a strong systematic bias towards selecting “large” cells as neighbors (i.e., cells with a large total UMI count), resulting in smoothed cells that on average contained a much larger UMI count than the cluster profile that was used as the basis for the simulation of these cells. Since the first step of kNN-smoothing is identical to that of one-step smoothing with *k*=1, it shared this bias for large cells in its first step. Astonishingly, the opposite was true for neighbors selected in its second step (*k* = 3), when smoothed cells exhibited smaller-than-average sizes. However, by the fourth step (*k* = 15), the average sizes were very close to the true cluster values in both datasets. The log-TPM variant exhibited similar behavior, but the distribution of sizes was generally much more spread out. Based on theoretical considerations, we think that it is undesirable for an algorithm to exhibit an overly strong size bias, as it will make very uneven use of the information available (see Discussion). We therefore believe that the near-convergence of the average cell size to the true cluster UMI count, as achieved by the kNN-smoothing algorithm for *k* ≥ 15, represents a desirable property that again makes kNN-smoothing preferable to the algorithm variants examined. In summary, our evaluation of the effects of our initial design decisions validated those decisions, as they resulted in an algorithm that provides more accurate results, and makes more even use of information from cells that differ in their total UMI counts (e.g., due to efficiency noise).

### The kNN-smoothing algorithm fails to accurately identify neighbors in scRNA-Seq data containing distinct T cell subsets

In the results presented above, the kNN-smoothing algorithm was applied to datasets comprising cell populations with very different expression profiles. For example, PBMCs consist mostly of monocytes, T cells, and B cells, and each of these cell types exhibits an expression profile that is very distinct from those of all the others. To obtain scRNA-Seq data for cells with known identities and very similar expression profiles, we downloaded data from various subsets of T cells, bead-enriched from human PBMCs (Zheng et al., 2017). On average, each T cell expression profile had approx. 1,500 transcripts, 50% of which belonged to genes encoding ribosomal proteins (see Figure S6a). When we combined all T cell datasets (see Methods) and performed PCA, we noticed that the first two PCs appeared to represent differences in ribosomal gene expression levels (see Figure S6b). To guard against the possibility that these differences represented batch effects rather than genuine biological differences, we decided to remove ribosomal genes from the data. A PCA on the remaining data no longer displayed obvious batch effects. However, the first PC still appeared correlated with the ribosomal gene content in the original data (see Figure S6c), suggesting that perhaps different T cell subsets exhibit differences in ribosome content.

To test the ability of kNN-smoothing to accurately identify neighbors in datasets containing cells from populations with very similar expression profiles, we combined expression profiles from the downloaded datasets to create three artificial datasets (see Methods). Each artificial dataset consisted of 1,000 profiles from naive CD4 T cells and 1,000 profiles from a different population, namely naive CD8 T cells (the first dataset), memory CD4 T cells (the second dataset), and B cells (the third dataset, serving as a control). In terms of their transcriptome, naive CD4 T cells were more similar to naive CD8 T cells than to memory CD4 T cells, however all three T cell subsets were much more similar to each other than to B cells (see Figure S7a). We first performed PCA on the unsmoothed data, which showed that the first PC perfectly separated the two cell populations for the B cell dataset (see Figure S7f, left), but not for the other two datasets(see Figure 8a, left, and Figure S7c, left), again highlighting that B cells were much more easily distinguishable from naive CD4 T cells than either naive CD8 T cells or memory CD4 T cells. In particular, the first two PCs captured only 2.7% and 1.4% of the total variance in the data, respectively, for the naive CD8 and memory CD4 T cell datasets, suggesting that technical noise, rather than the difference between the two cell populations, was the dominant source of variance. We then applied kNN-smoothing to each dataset, expecting that the smoothed expression profiles from the two populations were more clearly separated in principal component space than the unsmoothed profiles. However, for the naive CD8 and memory CD4 T cell datasets, smoothing led to a blurring, rather than a separation, of profiles from the different T cell subsets (see Figure 8a, center, and Figure S7c, center), demonstrating that the algorithm failed to consistently select neighbors from the same population when smoothing each expression profile. In contrast, the algorithm had no difficulties smoothing the expression profiles for the B cell dataset (see Figure S7f, center), suggesting that the problem originated from the fact that the various T cell subsets exhibited very similar expression profiles.

**Figure 8.**
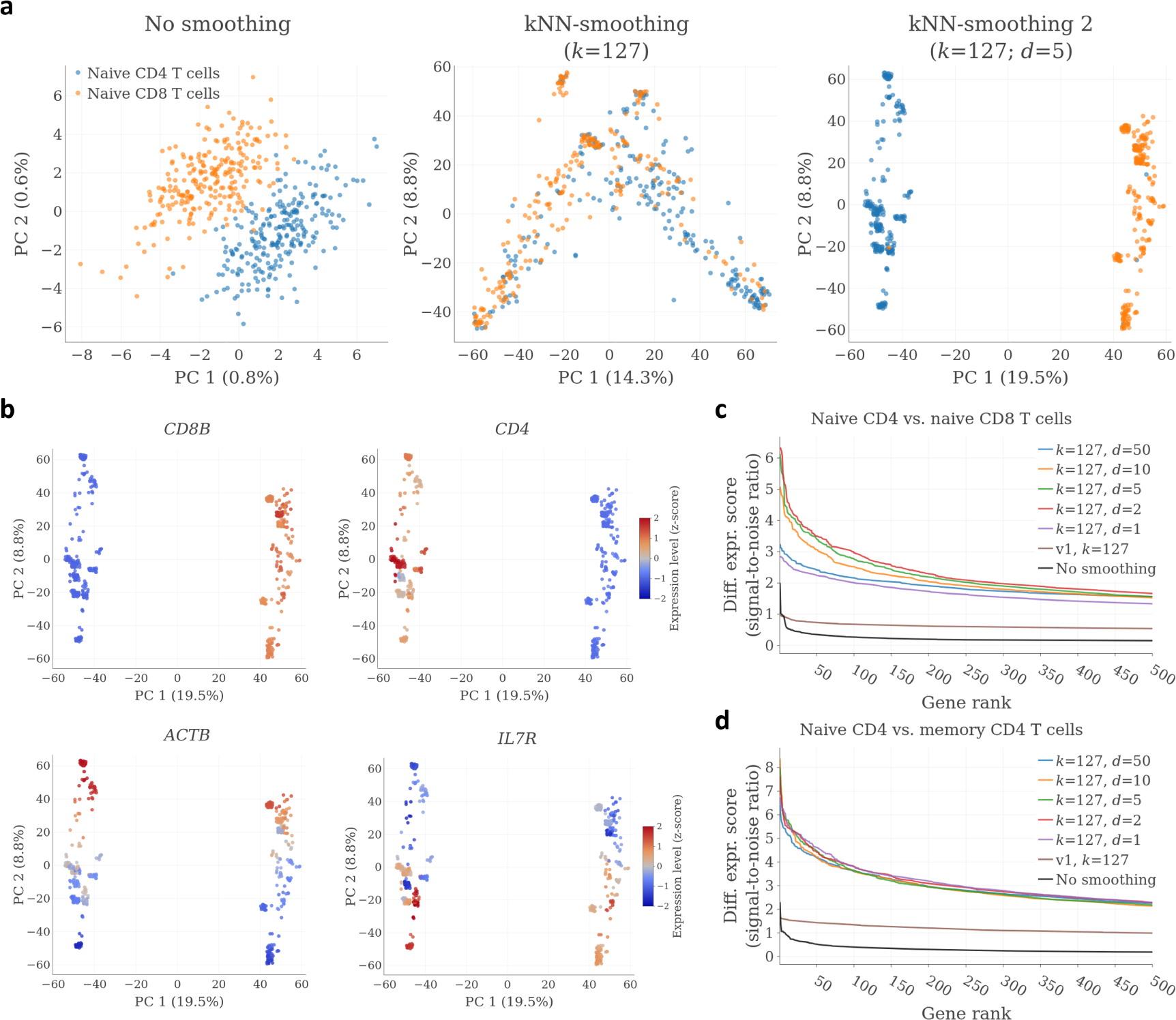
Accuracy of kNN-smoothing 2 for datasets containing subsets of T cells. **a** PCA plot for a dataset consisting of 1,000 profiles each from naive CD4 and CD8 T cells, respectively. Left, before smoothing; middle, after smoothing with kNN-smoothing; right, after smoothing with kNN-smoothing 2. A random subset of 250 cells from each population are shown to improve the readability of the figure. **b** Expression levels of four genes overlaid on a PCA plot of the cells after smoothing with kNN-smoothing 2. **c** Quantitative analysis of the smoothing accuracy for the CD4/CD8 T cell data using a differential expression metric. Shown are the differential expression scores, ranked from high to low, after smoothing with different parameters. **d** Same analysis as in (**c**), but for a dataset consisting of naive CD4 and memory CD4 T cells.

### An improved smoothing algorithm accurately identifies neighbors in scRNA-Seq data containing distinct T cell subsets

For closely related cell types, such as naive CD4 and CD8 T cells, it is reasonable to assume that most expressed genes are expressed at identical or near-identical levels in both cell types. Therefore, when datasets are composed of cells from either type, most expressed genes contain little or no information to help establish an accurate set of nearest neighbors for each cell. At the same time, those genes still contribute technical noise, which can drown out the expression differences of the small set of genes that are truly differentially expressed. According to this logic, it was not surprising that kNN-smoothing failed to correctly identify neighbors for datasets containing different subsets of T cells (see above). Since we had observed that PCA was nevertheless able to partially separate the cell populations, we reasoned that it would be possible to more accurately identify neighbors in such scenarios by calculating distances in principal component space. We therefore modified the kNN-smoothing algorithm so that in each step, the partially smoothed expression data are projected onto the first *d* principal components before Euclidean distances are calculated. The resulting algorithm, which we refer to as “kNN-smoothing 2”, is formalized in Algorithm 2.

We applied kNN-smoothing 2 to the combined T cell datasets (with *k*=127 and *d*=5), as before, and found that it resulted in very well-separated populations of cells (see Figure 8a, right, and Figure S7c, right). For both the naive CD8 T cell and the memory CD4 T cell dataset, cells were perfectly separated by type along the first principal component, with the exception of a few outlier cells that appeared to belong to the wrong cluster. As these cells did not exhibit an intermediate expression profile after smoothing, it appeared likely that these cells represented genuine contaminants in the bead-enriched samples, rather than smoothing artifacts. More importantly, the first PC now captured a substantial fraction of total variance in both datasets (19.5% and 30.9%, respectively), indicating that smoothing significantly improved the signal-to-noise ratio of the data. For the CD8 T cell dataset, we next examined the expression patterns of *CD8B* and and *CD4*, and found that their smoothed expression levels perfectly matched the expected patterns (see Figure 8b, top row). We also noticed that cells from both populations exhibited heterogeneity with respect to the second PC, which captured 8.8% of total variance. Among the genes with the strongest contribution to PC 2 were *ACTB* (beta-actin) and *IL7R* (CD127), exhibiting anti-correlated expression patterns (see Figure 8b, bottom row). As the actin cytoskeleton and *IL7R* are known to play important roles in TCR signaling and T cell homeostasis, respectively (Kumari et al. 2014; Carrette and Surh 2012), the observed expression heterogeneity could reflect different activation states of naive T cells in peripheral blood. We also observed heterogeneity in a number of immediate early genes (e.g., *IEG2* and *JUN*; see Figure S7b), which was recently reported as a common expression artifact associated with single-cell sample preparation (Brink et al. 2017). The biological importance of the observed expression heterogeneity was therefore unclear, and warrants further investigation. The smoothing results for the B cell dataset were not substantially different from those obtained with the earlier version of the kNN-smoothing algorithm (see Figure S7f, right).

As the kNN-smoothing 2 algorithm introduces an additional parameter, *d*, we next aimed to examine how different choices of *d* affected the smoothing results. We tested settings of *d* ranging from 1 to 50 on all three datasets, and compared results by ranking genes by their differential expression scores, which we defined to quantify the difference in expression of a gene in the two cell populations, relative to the technical noise (see Methods). We found that for the naive CD8 T cell dataset, kNN-smoothing 2 gave similar performance for settings of *d* ranging from 2 to 10 (see Figure 8c), whereas for the memory CD4 T cell dataset, results were stable across the entire range of values tested (see Figure 8d), perhaps reflecting the fact that distinguishing between naive and memory T cells was less challenging than distinguishing between naive CD4 and CD8 T cells. For both datasets, kNN-smoothing 2 clearly outperformed the first version of the kNN-smoothing algorithm for all settings of *d*. However, for the third dataset, containing naive CD4 T cells and B cells, the performance differences appeared negligible (see Figure S7e), again highlighting that kNN-smoothing 2 specifically improved smoothing accuracy for datasets containing highly similar cell types.

#### Algorithm 2 K-nearest neighbor smoothing 2 for UMI-filtered scRNA-Seq data

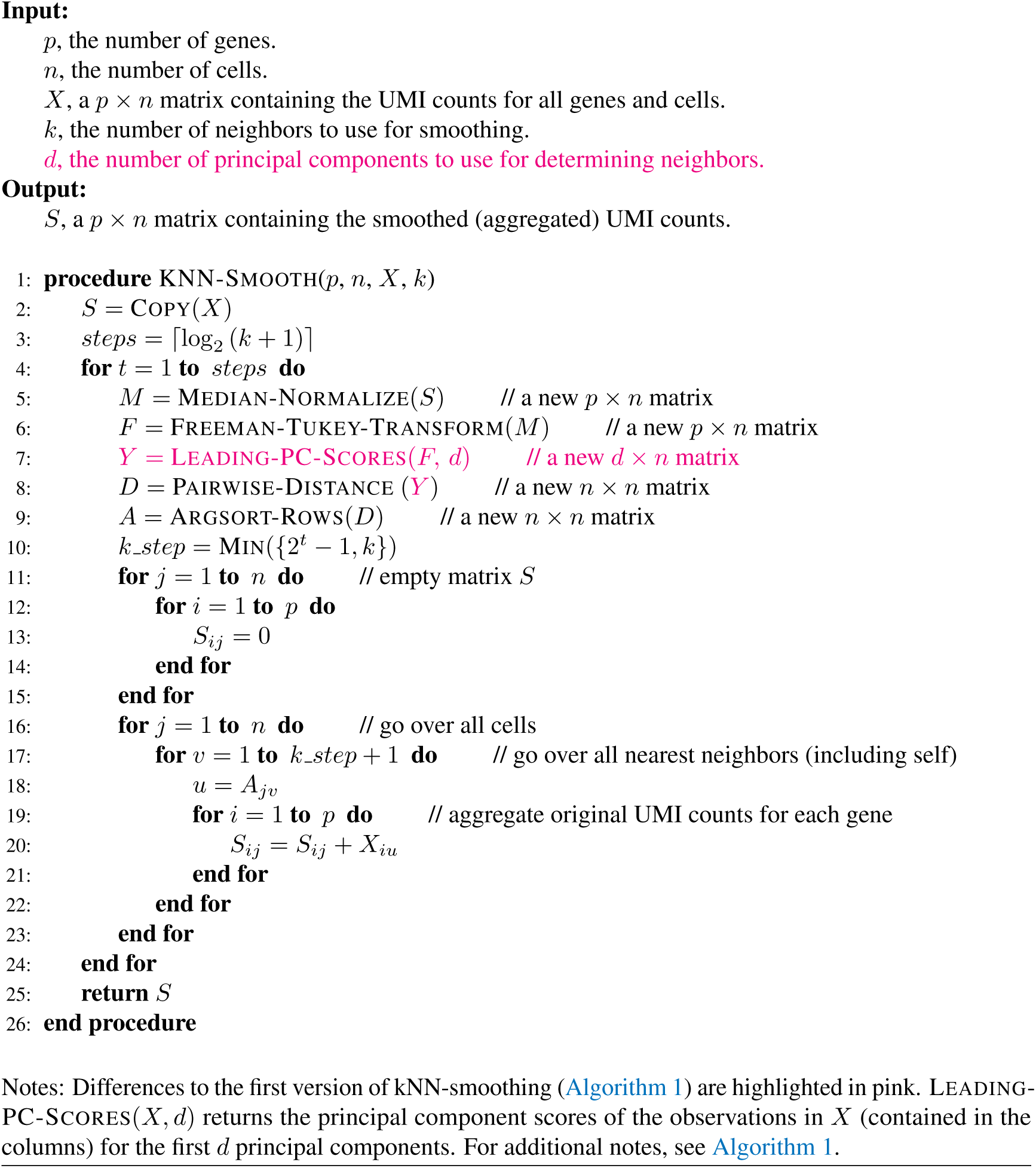

### kNN-smoothing 2 facilitates the identification of T cell subsets in peripheral blood

As different subsets of T cells generally exhibit very similar transcriptomes and contain relatively little mRNA (see above), their precise identification in scRNA-Seq data is a challenging task. In previous reports, authors have struggled to delineate different T cell subsets in peripheral blood using clustering (see Figure 3j in Zheng et al. (2017)), and to distinguish regulatory T cells from other T cells found in mouse spleens (Zemmour et al. 2018). To test whether kNN-smoothing 2 would facilitate the identification of T cell subsets found in peripheral blood, we downloaded a 10x Genomics pan-T cell dataset containing 4,583 expression profiles, applied kNN-smoothing 2, and generated expression heatmaps showing the expression patterns of the most highly variable genes following hierarchical clustering (see Figure 9a, top). We next aimed to determine, for each expression profile in the data, whether it represented a naive or a memory T cell. To do so, we first determined a set of marker genes for naive and memory T cells, by identifying genes with differential expression between experimentally isolated naive and memory T cell subsets (see Figure S8a and Methods). We examined the expression patterns of these marker genes in the smoothed and clustered pan-T cell data, and found that naive and memory marker genes exhibited mutually exclusive expression patterns, and that there was a very clear clustering of cells expressing either set of marker genes (see Figure 9a, middle). Finally, we also aimed to compare the smoothed expression profiles to subset-specific expression profiles, again obtained from experimentally isolated T cell subsets (see Methods). We found that the cells expressing naive and memory marker genes also had expression profiles that were specifically correlated with those of naive and memory T cell subsets, respectively (see Methods). A small cluster of cells had profiles that partly resembled naive T cells, but also expressed a set of genes not generally observed in those cells. This signature partially overlapped with the set of genes associated with heterogeneity among naive T cells observed earlier (see Figure S7b), including *CD7*, *CD3D*, and *GZMM*. The possibility that this cluster of cells represents a specific activation state of naive T cells requires further investigation. In summary, these results demonstrate that kNN-smoothing 2 enabled the identification of naive and memory T cells in pan T-cell scRNA-Seq data.

**Figure 9.**
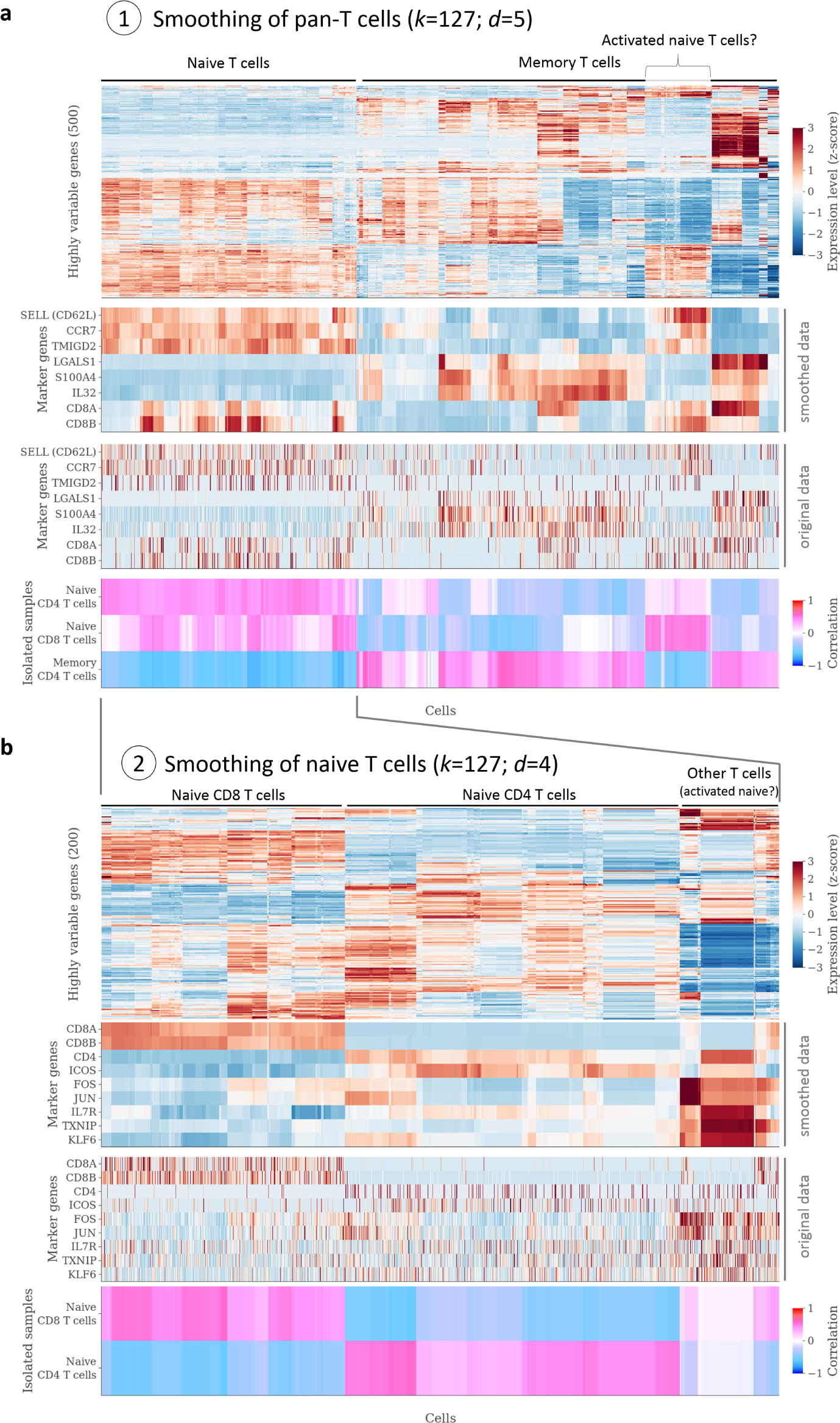
Identification of T cell subsets in a peripheral blood pan-T cell data using kNN-smoothing 2. **aa** Analysis of 4,583 pan-T cell expression profiles. Top, heatmap of hierarchically clustered data after application of kNN-smoothing 2 with *k*=127 and *d*=5. Below, expression of naive and memory T cell marker genes in the smoothed and unsmoothed data, using the same ordering of cells as in the heatmap above. Bottom, correlation with expression profiles from isolated subsets of naive and memory T cells, using the same ordering of cells as in the heatmap above (see Methods for details). **b** Analysis of a subset of 1,656 profiles identified as naive T cells, using kNN-smoothing with *k*=127 and *d*=4. Panels are organized as in **a**.

To further test the ability of kNN-smoothing 2 to distinguish T cell subsets in heterogeneous data, we repeated the analysis (smoothing, hierarchical clustering, examination of marker genes, and correlation with expression profiles from experimentally isolated subsets) on the group of 1,656 cells that were identified as naive T cells in the previous step, hoping that we would be able to distinguish naive CD4 from naive CD8 T cells (see Figure 9b and Figure S8b). We found that based on the expression of marker genes and the correlation analysis, we were indeed able to distinguish these subsets, with CD8 T cells exhibiting stable expression of *CD8A* and *CD8B* after smoothing. Moreover, smoothing was able to recover the expression of *CD4* in CD4 T cells, even though this gene appeared to expressed at much lower levels compared to *CD8A* and *CD8B*. A small set of of cells again exhibited an “activation” expression signature which we found to be very similar to the that of the previously discussed “activated naive T cells” (data not shown). In summary, these results demonstrate that kNN-smoothing 2 enabled the identification of naive CD4 and naive CD8 T cells in pan T-cell scRNA-Seq data.

### Python implementations of kNN-smoothing and kNN-smoothing 2 process datasets containing thousands of cells within a few minutes

For a smoothing method to be of practical use, it not only needs to provide accurate results, but it must also finish in a reasonable amount of time. We therefore measured runtimes of our Python implementations of kNN-smoothing and kNN-smoothing 2 on Chromium PBMC data containing 21,425 expressed genes, using subsampling to generate datasets with sizes ranging from *n*=2,000 to *n*=8,000 cells, on a laptop with an Intel^®^ Core™ i7-6600U processor and 20 GiB of memory (see Methods). We found that the runtimes ranged from a few seconds to just over 14 minutes (for *k*=511 and *n*=8,000), and that runtime increased linearly with *k* (see Figure 10a). For *n*=2,000 and *n*=4,000, both algorithms had very similar runtimes. However, for *n*=8,000, kNN-smoothing 2 was slightly faster than kNN-smoothing, because the calculation of pairwise distances was significantly faster when the data had been reduced to *d* dimensions (corresponding to the first *d* PCs), and because our implementation relied on an efficient algorithm for calculating the first *d* PCs (Halko, Martinsson, and Tropp 2009).

**Figure 10.**
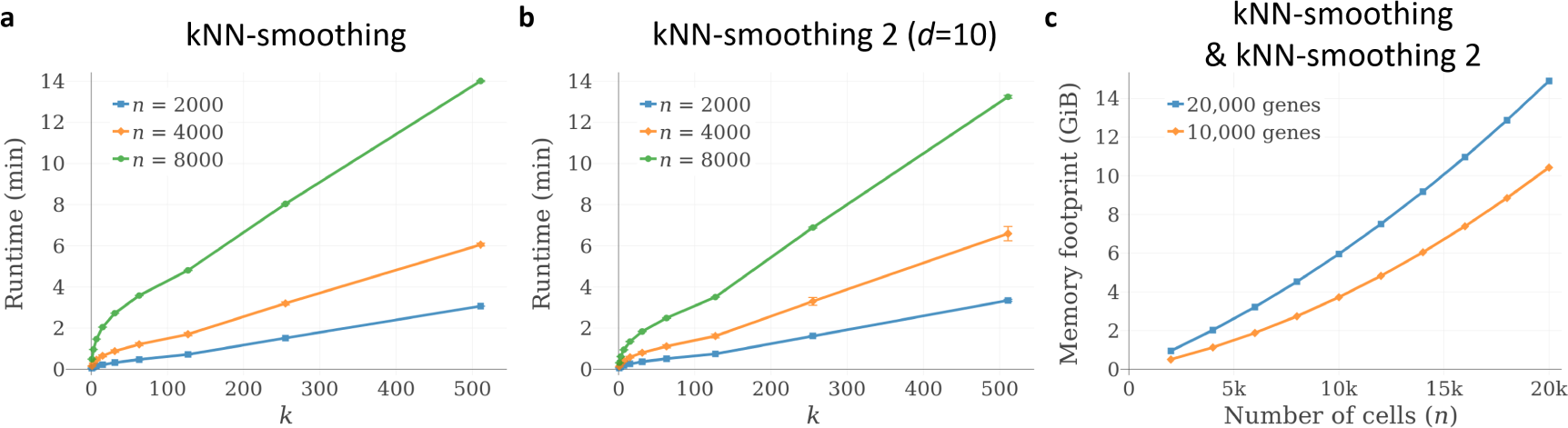
Performance and memory footprint of kNN-smoothing and kNN-smoothing 2 for datasets of different sizes. **a, b** Runtimes of Python implementations of the kNN-smoothing and kNN-smoothing 2 algorithms, respectively, when applied to datasets obtained by subsampling different numbers of cells (*n*) from a scRNA-Seq dataset of human peripheral blood mononuclear cells (PBMCs), published online by 10x Genomics. Smoothing was performed on 21,415 genes with expression. Settings of *k* are indicated on the x-axes. **c** Predicted memory footprint of the kNN-smoothing algorithms as a function of the number of cells in the dataset (*n*). See Methods for details.

We also calculated the memory footprint of our Python implementation of kNN-smoothing, which requires three copies of the expression matrix (original, smoothed, smoothed and transformed) and two *n*-by-*n* arrays (the distance matrix and a sorted indexing array) to be held in memory. We assumed that each expression measurement would be represented in memory by an 8-byte floating point value. From the results Figure 10b, it appears that for datasets containing approx. 20,000 protein-coding genes, the largest datasets that can be analyzed (without memory swapping) contain approx. 5k, 10k, and 20k cells, for computers with 4 GiB, 8 GiB, and 16 GiB of memory, respectively. Overall, these results demonstrate that kNN-smoothing can be run on most laptops and PCs for datasets containing several thousand cells, in a time-span of minutes or even seconds. The memory requirements for kNN-smoothing 2 are not substantially different from that of kNN-smoothing, as the storage of the first *d* principal component scores requires relatively little additional memory.

## DISCUSSION

### Design and applicability of kNN-smoothing and kNN-smoothing 2

In this work, we have proposed *k-nearest neighbor smoothing* (kNN-smoothing), and kNN-smoothing 2, two novel algorithms for smoothing high-throughput scRNA-Seq data, aimed at significantly improving the signal-to-noise ratios of the gene expression values for each cell by aggregating information from similar cells (“neighbors”). The first innovation of kNN-smoothing is the step-wise approach to identifying the nearest neighbors, which we have shown to result in more accurate results compared to a one-step appraoch. The second innovation is the use of the Freeman-Tukey transform, designed to ensure that the expression levels of all genes contribute approximately equally in the distance calculation, independent of their general expression level. kNN-smoothing 2 combines these innovations with principal component analysis, which helps to filter out noise and focus on the most salient expression differences among cells. kNN-smoothing 2 is more efficient, powerful, and flexible relative to the first version of kNN-smoothing, and should therefore be the preferred choice for practically all applications.

It might appear that by smoothing single-cell data, one is compromising on important information pertaining to the individuality of each cell. We note that while cell-to-cell variation within a given cell type is of clear importance, in most applications one is querying for cell populations that are each represented by an appreciable number of cells. Thus, given the routine profiling of thousands or even tens of thousands of cells, and the inherent noisiness of the data under study, our smoothing algorithm offers a clear advantage in terms of the identification of those populations.

We designed the kNN-smoothing algorithm based on the observation that data from multiple high-throughput scRNA-Seq protocols (including inDrop, Drop-seq, and 10x Genomics’ Chromium) share common technical noise characteristics. Specifically, after the application of “median-normalization” to account for efficiency noise, the gene expression values in technical replicates are approximately Poisson-distributed. We believe that this is a direct consequence of the fact that all of these protocols only capture a small fraction of transcripts of each cell, employ 3’-or 5’-end counting (“tagging”), and avoid overcounting of amplified transcripts by UMI-filtering. Therefore, we predict that the Poisson noise characteristic applies to all such scRNA-Seq protocols that use UMI filtering, but not to other scRNA-Seq protocols. This idea clearly warrants a more detailed investigation, which is beyond the scope of this paper. Whatever the origins of the noise characteristics described here, the fact that they are shared between the aforementioned protocols implies that our proposed algorithms are in principle applicable to any dataset generated using those protocols.

### Comparison with previously reported methods

Our algorithm combines a previously proposed normalization method (Grün, Kester, and Oudenaarden 2014) with a standard variance-stabilizing transformation (VST) for Poisson-distributed data (Freeman and Tukey 1950). We are not aware of prior work suggesting the use of a VST in the context of smoothing scRNA-Seq data. Instead, most work has focused on parametric modeling (see Introduction). While these approaches can certainly be effective, our work suggests that they are not strictly necessary to effectively to address the issue of noise in scRNA-Seq data. Moreover, sophisticated models often require complex inference procedures, which can be difficult to implement correctly and efficiently. In contrast, our method requires only a few lines of code, while still being based on statistical theory, and our Python implementation runs in a matter of seconds or minutes on datasets containing a few thousand cells.

Simple aggregation or averaging of scRNA-Seq expression profiles has been previously employed in specific contexts, for example for library size normalization (Lun, Bach, and Marioni 2016). Recently, La Manno et al. (2017) employed a simple version of k-nearest neighbor smoothing (“pooling”) as part of a method designed to estimate the time derivative of mRNA abundance based on unspliced RNA sequences. The authors defined the most similar cells based on log-transformed data (for read counts from the SMART-Seq2 protocol), or PCA-transformed data (for UMI counts from inDrop and 10x Genomics protocols). However, they did not provide any justification for their choices of similarity metrics, a discussion of the statistical properties of the data before and after smoothing, or a quantification of the gain in expression accuracies achieved. Moreover, neither of these studies aimed to develop a general-purpose method to improve the signal-to-noise ratio of scRNA-Seq data, or employed a step-wise approach for defining the nearest neighbors, as we have done here. Our work can be compared to other recently proposed methods that aim to specifically address the issue of technical noise in scRNA-Seq data: Dijk et al. (2017) aimed to apply the idea of manifold learning using diffusion maps to scRNA-Seq data (see Supplementary Text for a demonstration of kNN-smoothing on one of the datasets analyzed in their study), and W. V. Li and J. J. Li 2017) generated Poisson-distributed expression data. Dijk et al. (2017) started from bulk microarray expression data, which was then “downsampled using an exponential distribution” to obtain specific proportions of zero values, while W. V. Li and J. J. Li (2017) defined gene-specific “dropout rate[s]”, and set individual expression values to zero using Bernoulli trials with those rates. Based on the results presented in this work, we believe that neither of these approaches faithfully reproduces the noise characteristics of UMI-filtered scRNA-Seq data.

### Use of simulation studies to quantify the accuracy of scRNA-Seq smoothing methods

As scRNA-Seq is currently the only technology that can be used to interrogate complete transcriptomes of single cells in a highly parallelized fashion, there exist no “gold standard” datasets to benchmark scRNA-Seq smoothing algorithms (i.e., datasets that contain a heterogeneous mixture of cells whose true single-cell expression profiles have been determined using an orthogonal method). Therefore, one most resort to simulation studies in order to quantitatively assess the accuracies of smoothing methods. Here, we established a new method for using real scRNA-Seq datasets to simulate UMI-filtered scRNA-Seq data that consist of a mixture of cell types (clusters). The simulated data exhibit Poisson-distributed sampling noise, modeled on top of efficiency noise, for which we used the observed distribution of total UMI counts per cell in the real data. (This might result in an overestimate of efficiency noise, as some differences in total UMI counts could also reflect biological differences in total mRNA abundance and/or cell size.) Our methodology is based on the understanding of the sources and characteristics of technical noise in UMI-filtered scRNA-Seq data as described in this work, and a visual comparison between the real and the synthetic datasets led us to conclude that it can also reproduce the majority of the biological heterogeneity observed in the real dataset. For the analyses reported here, we decided to limit the simulations to *K* =10 clusters, but the procedure is compatible with any integer choice of *K* for 1 ≤ *K* ≤ *n* (where *n* is the number of cells in the real data), and the use of hierarchical clustering ensures consistency between datasets generated using similar choices of *K* (e.g., for *K* = 11, one of the clusters present in the *K* = 10 dataset would be split into two distinct clusters, while all other clusters remain identical).

Based on the simulated data, we were able to show that with *k* ≥ 7, kNN-smoothing produced much more accurate results for both simulated datasets, when compared to MAGIC (Dijk et al. 2017) and scImpute (W. V. Li and J. J. Li 2017). This was true for all MAGIC and scImpute parameter settings tested, independently of whether we quantified accuracy using both relative (PCC) or absolute (RMSE) measures, and independently of whether we used log_2_-transformed or square root-transformed expression values in these calculations. In some cases, kNN-smoothing was able to recover the true expression profile with near-perfect accuracy, which we never observed for either of the two other methods. Our results therefore suggest that kNN-smoothing generally outperforms MAGIC and scImpute on UMI-filtered scRNA-Seq data containing highly heterogeneous cell populations.

A limitation of our approach to simulating scRNA-Seq data is that it ignores certain biological sources of heterogeneity: For example, cells from the same cell type might be in different cell cycle phases, and these differences would be lost (averaged out) as part of the simulation procedure. More generally, our current approach is unable to simulate datasets that contain a mixture of cells from different stages of a continuous dynamic process (such as cell differentiation), and procedures that can simulate UMI-filtered scRNA-Seq data for those types of experiments need to be established in order to quantitatively evaluate the performance of smoothing methods in such scenarios.

### How to choose *k* and *d*?

kNN-smoothing depends on one parameter (*k*), whereas kNN-smoothing 2 depends on two parameters (*k* and *d*). In addressing the question of how to choose these parameters, we would like to emphasize that kNN-smoothing was primarily designed as a tool to facilitate exploratory data analysis (EDA). In EDA, the primary goal is to learn about the relevant aspects of the data (e.g., the set of cell populations present, the), using whatever means conducive to this goal. In this context, there exist no objectively “correct” or “incorrect” parameter settings. Rather, we are interested in establishing guidelines for choosing settings that provide the greatest insight into the data at hand. In practice, we generally encourage the testing of different parameter settings and a comparison of the different results obtained.

The results obtained when applying kNN-smoothing or kNN-smoothing 2 to a particular dataset strongly depend on the choice of *k*. Choosing *k* very small might not adequately reduce noise. On the other hand, choosing *k* too large incurs the risk of smoothing over biologically relevant expression heterogeneity, and can also lead to artifactual expression profiles that consist of averages of profiles belonging to different cell populations. As a result, our method provides no guarantee that smoothed expression profiles represent existing cell populations.

In exploring data from heterogeneous tissues, it seems clear that the choice of *k* should be guided by the estimated number of cells present from each population. However, different populations can be present at very different abundances: As an extreme example, imagine that in a dataset containing 1000 cells, 900 cells belong to population A, 90 cells belong to population B, and 10 cells belong to population C. A conservative approach would then be to apply kNN-smoothing with *k*=9, thereby avoiding any “over-smoothing”. However, it is possible that this degree of smoothing is insufficient to reliably distinguish between cells from populations A and B, when in fact one could choose *k*=89 without over-smoothing cells from either population. To avoid underutilizing the data available, we therefore find it advisable to smooth more aggressively first and to identify the main clusters apparent in the data. Then, for each cluster separately, smoothing with smaller *k* can be re-applied to the original, unsmoothed data, in order to identify any subpopulations that had perhaps been over-smoothed in the previous step. We think that such a hierarchical strategy to identifying cell populations is broadly applicable and significantly accelerates the analysis of heterogeneous data with highly variable population abundances.

An appropriate choice of *k* also depends on the particular application: When analyzing cells undergoing a highly dynamic process (e.g., differentiation), large values of *k* might result in an overly coarse picture of the transcriptomic changes. In contrast, when aiming to distinguish distinct cell types, larger choices of *k* can help identify robust expression profiles for each type.

Finally, our analyses suggest that the choice of *d* is generally less critical than that of *k*, as different choices often gave very similar results. However, overly small or large values of *d* can lead to inaccurate smoothing results. For highly heterogeneous datasets, setting *d* too small can lead to a loss of resolution, as biological differences captured by the higher PCs are ignored. For less heterogeneous data, setting *d* very large can lead to poor results, as many PCs only capture technical noise and drown out the biological differences captured by the first PCs. We propose a default setting of *d*=10 for applications to scRNA-Seq data from heterogeneous tissues. One strategy is to identify a suitable setting of *k* using *d*=10, and to then test if changing *d* can improve the results. We think that as in other applications involving PCA, the percentage of total variance explained by the first *d* components (in the smoothed data, after FT-transform) can serve as a guide to selecting *d*. However, we hesitate to recommend a specific percentage (e.g., 80%), as the fraction of variance that represents biological differences depends on the degree of smoothing as well as the heterogeneity in the data. Generally speaking, we recommend to err on the side of including too many PCs, in order to avoid the loss of biological signals.

### Importance of smoothing for the analysis of scRNA-Seq data

We have demonstrated the application of kNN-smoothing to data generated using the inDrop (Klein et al. 2015) and Chromium (Zheng et al. 2017) protocols, and shown that in both cases, the algorithm was able to recover cell type-specific expression patterns for previously described marker genes. Moreover, the achieved noise reduction made it straightforward to apply hierarchical clustering (Eisen et al. 1998), a powerful method for exploratory analysis of gene expression data that performs poorly on unsmoothed scRNA-seq data. We obtained similar results when we applied kNN-smoothing 2 to peripheral blood pan-T cell data, where we were able to demonstrate that smoothing made it possible to discriminate between naive and memory T cells, and also between naive CD4 and naive CD8 T cells, using purely unsupervised methods.

kNN-smoothing, specifically kNN-smoothing 2, has the potential to improve the performance of many advanced analysis methods that rely on PCA or other dimensionality reduction techniques, including methods for trajectory inference (e.g., Cao et al. 2017). Importantly, kNN-smoothing works by aggregating information across cells, rather than across genes. Therefore, it avoids the introduction of artificial gene-gene dependencies, which are highly problematic when downstream analyses involve methods whose null models assume independence between genes, such as GO enrichment analysis (Subramanian et al. 2005; Eden et al. 2009). At the same time, kNN-smoothing clearly introduces dependencies between cells. Naturally, the extent to which this is the case depends on the magnitude of *k*.

Recently, researchers and funding bodies have proposed the generation of “cell atlases”, systematic efforts aimed at providing exhaustive molecular descriptions of all cell types and states present in human tissues under healthy as well as disease conditions such as cancer (Regev et al. 2017; National Cancer Institute 2017; The Chan Zuckerberg Initiative 2018). As scRNA-Seq is generally seen as an important experimental tool for the realization of these projects, kNN-smoothing 2 could represent a valuable analysis tool for the identification of novel cell types and states, as well as for the characterization of their expression profiles.

### Implications for study design

We have shown that there exists a quadratic relationship between “cell coverage” (the number of profiles obtained for a given population of cells) and the potential accuracy with which the average expression levels of individual genes in cells from that population can be quantified. To improve the signal-to-noise ratio by a factor of two, the cell coverage needs to be quadrupled. In studying heterogeneous tissues, researchers should therefore consider the estimated frequencies of the different populations of interest, and adjust the number of cells profiled accordingly. For example, if a population comprising only 1% of cells is to be “covered” by 20 profiles (on average), this implies that 2,000 cells need to be profiled. Using simple binomial statistics, approx. 2,800 cells would then have to be profiled for a 95% chance of obtaining at least 20 profiles from that population.

In addition, the relationship between cell coverage and quantification accuracy brings into focus the question of what constitutes an optimal number of sequencing reads per cell. While a quantitative treatment of this issue is beyond the scope of this work, it is clear that in many situations, it would be more beneficial to sequence additional cells, rather than increase the read coverage per cell. The precise optimum likely depends on numerous factors, and is difficult to determine without an examination of all the experimental, statistical, and computational factors involved in scRNA-Seq studies. However, since sequencing can represent the single most expensive part of the experiment, this issue clearly warrants further investigation.

Based on the work described here, it is tempting to speculate that in theory, there is no limit as to how accurately the average expression profile of individual cell populations and sub-populations can be determined using scRNA-Seq. Our analysis suggests that the signal-to-noise ratio can always be improved by aggregating more profiles from “biologically identical” cells. In practice, however, the number of cells that can be analyzed is limited by the protocol used, the cost of the experiment, the number of cells available, and/or the rarity of the population of interest. Furthermore, the accuracy with which “biologically identical” cells can be identified based on their noisy profiles depends on several factors, including the level of granularity required and the number of transcripts present in the cell. It is therefore not clear whether all biologically relevant cell types and states can be accurately identified with current scRNA-Seq protocols.

### Conclusions

In this work, we have used multiple datasets to demonstrate that PCA and hierarchical clustering, two basic techniques for analyzing gene expression data benefit strongly from kNN-smoothing. In future work, it would be interesting to explore the effect of smoothing for additional types of analyses, including differential expression analysis, gene set enrichment analysis, and trajectory inference. We anticipate that our kNN-smoothing 2 algorithm will benefit all of these approaches, and generally enable the more effective analysis of scRNA-Seq data in wide variety of settings. It should again be noted, however, that smoothed expression profiles of cells are no longer statistically independent, so smoothing should not be used naively in combination with statistical tests for differential expression.

Given the rapidly increasing number of cells that can be profiled in a single experiment (Svensson, Vento-Tormo, and Teichmann 2018), an important direction for future research is how to make the kNN-smoothing 2 algorithm scalable to datasets containing tens or hundreds of thousands of cells. In these instances, calculating a matrix containing all pairwise distances can exceed memory limitations, and smoothing can become prohibitively slow. A strategy to dealing with these situations could involve downsampling of cells, combined with a hierarchical approach to smoothing as outlined above.

High-throughput scRNA-Seq technologies are widely believed to hold enormous potential for studying heterogeneous tissues and dynamic cellular processes in health and disease. However, the inherent noisiness of the data require that in parallel to advances in experimental technologies, innovations in the areas of algorithm development and machine learning are necessary in order to realize this potential. Fortunately, data from different protocols exhibit very similar statistical properties, presumably due to their shared reliance on 3’-end counting and UMI filtering. These properties should directly inform the design of effective algorithms for the smoothing and analysis of scRNA-Seq data. We have described two closely related algorithms for smoothing scRNA-Seq data that are generally applicable, efficient, and easy-to-implement. The large improvements in the signal-to-noise ratio that can be achieved with these algorithms helps to expand the realm of possibilities for downstream analyses, and to better leverage scRNA-Seq for the understanding of complex biological systems.

## METHODS

### Download and processing of inDrop pure RNA replicate data

Raw sequencing data were downloaded from SRA (experiment accession SRX863258). In this experiment by Klein et al. (2015), droplets containing pure RNA extracted from K562 cells were processed using the inDrop protocol. The downloaded data were processed using a custom pipeline. Briefly, SRA data were converted to the FASTQ format using fastq-dump. Next, the “W1” adapter sequence of the inDrop RT primer were located in the barcode mate sequence (the first mate of the paired-end sequencing), by comparing the 22-mer sequences starting at positions 9-12 in the read with the known W1 sequence, allowing at most two mismatches. Reads for which the W1 sequence could not be located in this way were discarded. The start position of the W1 sequence was then used to infer the length of the first part of the inDrop cell barcode in each read, which can range from 8-11 bp, as well as the start position of the second part of the inDrop cell barcode, which always consists of 8 bp. Cell barcode sequences were mapped to the known list of 384 barcode sequences for each read, allowing at most one mismatch. The resulting barcode combination was used to identify the cell from which the read originated. Finally, the UMI sequence was extracted, and only with low-confidence base calls for the six bases comprising the UMI sequence (minimum PHRED score less than 20) were discarded. The mRNA mate sequences (the second mate of the paired-end-sequencing) were mapped to the human genome, release GRCh38, using STAR 2.5.3a with parameter “–outSAMmultNmax 1” and default parameters otherwise. Testing the overlap of mapped reads with exons of protein-coding genes and UMI-filtering was performed using custom Python scripts. Droplets (barcodes) were filtered for having a total UMI count of at least 10,000, resulting in a dataset containing UMI counts for 19,865 protein-coding genes across 935 droplets.

### Download of 10x Genomics ERCC spike-in expression data

UMI counts for ERCC spike-in RNA processed using the 10x Genomics scRNA-Seq protocol (Zheng et al. 2017) were downloaded from the 10x Genomic website. The dataset consisted of UMI counts for 92 spike-ins across 1,015 droplets.

### Download of Drop-Seq ERCC spike-in expression data

UMI counts for ERCC spike-in RNA processed using the 10x Genomics scRNA-Seq protocol (Macosko et al. 2015) were downloaded from GEO accession number GSM1629193. The dataset consisted of UMI counts for 80 spike-ins across 84 droplets.

### Prediction of scRNA-Seq noise characteristics based on Poisson statistics

In this paper, we initially focus on the technical variation observed in scRNA-Seq data for droplets containing identical pools of pure mRNA. Let 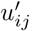 be the observed UMI count for the *i*’th gene (or ERCC spike-in) in the *j*’th droplet, for *i* = 1, …, *p* and *j* = 1, …, *n*. Similarly, let 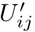 be a random variable representing the UMI count for the *i*’th gene in the *j*’th cell. We assume that 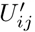 is Poisson-distributed with mean 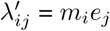, where *m*_*i*_ is the number of mRNA molecules present for the *i*’th gene, and *e*_*j*_ corresponding to the capture efficiency of the scRNA-Seq protocol for the *j*’th droplet (both *m*_*i*_ and *e*_*j*_ are unknown). We further assume that 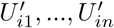 are independent, for all *i*. For the sake of simplicity, we assume that the read coverage (the number of reads sequenced per cell) is infinite, so that there are no cases in which a transcript is not observed due to limited read coverage. In practice, limited read coverage will not invalidate the Poisson assumption, but result in lower “effective” capture efficiencies.

If all *e*_*j*_ were identical (say, equal to *e*^global^), then 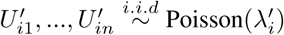, with 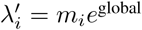. Grün, Kester, and Oudenaarden (2014) have proposed to normalize the expression profile of each cell to the median total UMI count across cells (Model I in Grün et al.), in order to counteract the differences in capture efficiency (“efficiency noise”). Median-normalization consists of calculating the total UMI count per profile (cell or droplet), 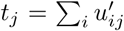, calculating the median *t*^med^ = median*{t*_1_*, …, t*_*n*_}, and then multiplying each 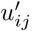 by the factor *t*^med^/*t*_*j*_.

Based on the results by Grün et al., we hypothesized that median-normalized data would be approximately Poisson-distributed, as long as the differences in capture efficiency were not too extreme. Therefore, we let 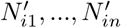 represent the UMI counts for the *i*’th gene after median-normalization, and assume them to be i.i.d. Poisson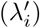.

For Poisson-distributed variables, the variance is always equal to the expectation (defined by *λ*). Let 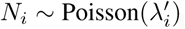. For the coefficient of variation (CV) of *N*_*i*_, we have:

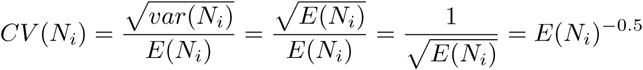

Taking the logarithm on both sides gives:

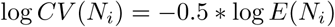

Therefore, the relationship between log *E*(*N*_*i*_) and log *CV* (*N*_*i*_) is linear with a slope of -0.5. This is indicated by the gray lines in Figure 1a-f.

The probability of observing a count of zero for *N*_*i*_ is given by the Poisson PMF:

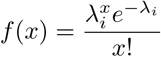

Therefore, *P* (*N*_*i*_ = 0) = *e*^−λ*i*^ values are shown as the orange lines in Figure 1g-i.

If a computational pipeline used to determine UMI counts reports systematically inflated values, then the median-normalized UMI counts for the *i*’th gene can be approximately represented by a scaled Poisson variable 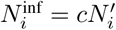, where *c* is the inflation factor. 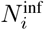 then has mean 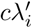 and variance 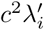 so for 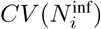, we have:

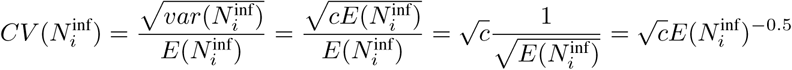

Taking the log on both sides gives:

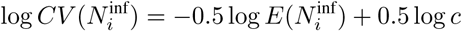

Therefore, the relationship between 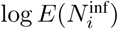 and 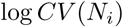 will still be linear, but with an y-axis intercept of 0.5 log *c* instead of 0, which is consistent with Figure 3b,e.

### Prediction of the effect of aggregating scRNA-Seq expression profiles from technical replicates

We again assume that for droplets containing identical pools of pure mRNA, the median-normalized UMI counts 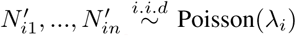. Let 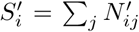, and 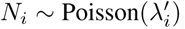. It is clear that 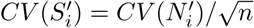:

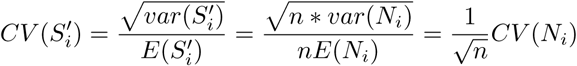

Similarly, for averaged UMI counts 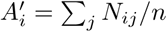:

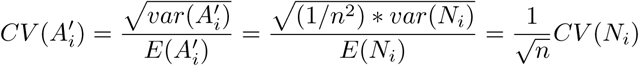

This effect is demonstrated in Figure 2.

### Smoothing of scRNA-Seq expression profiles from biological samples based on Poisson statistics

In real data, genes can exhibit differential expression across cells. Therefore, we define *λ*_*ij*_ = *m*_*ij*_*e*_*j*_, where *m*_*ij*_ is the number of mRNA molecules present for the *i*’th gene in the *j*’th cell, and *e*_*j*_ is the capture efficiency of the scRNA-Seq protocol for the *j*’th cell. Let *U*_*ij*_ be a random variable representing the UMI count for the *i*’th gene in the *j*’th cell. We again assume that *U*_*ij*_ is Poisson-distributed with mean *λ*_*ij*_, and that *U*_*i1*_,…*U*_*in*_ are independent, for all *i*. Let 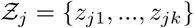 be the set of *k* nearest neighbors of the *j*’th cell, as determined in Algorithm 1. Let 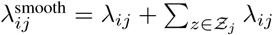. We then define the aggregated expression level 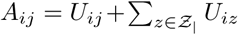, and note that 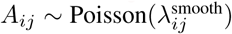. From the aforementioned discussion, it follows that if the *k* neighbors have transcriptomes that are sufficiently similar to that of the *j*’th cell, and if the efficiency noise is not too strong, then 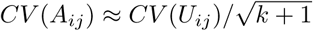. Similarly, we can calculate the averaged expression level *S*_*ij*_ = *A*_*ij*_/(*k* + 1). Then *S*_*ij*_ is a Poisson variable with mean 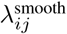, scaled by a factor of 1/(*k* + 1), and therefore has the same CV as *A*_*ij*_. The point here is that even if the *U*_*ij*_ are not identically distributed (due to expression differences and/or efficiency noise), simple aggregation or averaging will always result in Poisson-distributed smoothed values. The same is not true for weighted sums or averages. Let {*w*_*j*0_, *w*_*j*1_, …, *w*_*jk*_} represent weights (all positive), and let 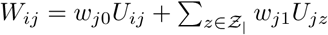. Then the weighted sum *W*_*ij*_ is neither a Poisson nor a scaled Poisson variable, unless all weights are identical.

### Download and processing of inDrop pancreatic islet data

Raw sequencing data were downloaded from SRA (experiment accession `SRX1935938`). In this experiment by Baron et al. (2016), inDrop was applied to pancreatic islet tissue from a human donor. Data was processed using the same pipeline used for the inDrop pure RNA data, and only profiles with a total UMI count of at least 1,000, resulting in a dataset containing UMI counts for 19,865 protein-coding genes across 2,109 cells. We refer to this dataset as the `PANCREAS` dataset.

### Download and processing of 10x Genomics Chromium (v2) peripheral blood mononuclear cell (PBMC) data

We downloaded the UMI-filtered expression matrix of the dataset titled “4k PBMCs from a Healthy Donor” from the 10x Genomics website (www.10xgenomics.com). The data was processed by 10x Genomics using the “Cell Ranger” software, version 2.1.0. A QC report of the dataset is available on the 10x Genomics website. The downloaded expression matrix contained 33,694 genes and 4,340 cells. We removed 13,921 genes that had no expression in the entire dataset, and then another 8 genes with duplicate gene names (keeping only the first instance of each gene). The final dataset contained 19,765 genes. We refer to this dataset as the `PMBC` dataset.

### Download and processing of mouse myeloid progenitor data

UMI counts were downloaded from GEO, accession number GSE72857. The 19 clusters for cells are available at `MAGIC`’s (Dijk et al. 2017) code repository: https://github.com/pkathail/magic/issues/34. 27,297 cells with cluster labels were used for performing k-nearest neighbor smoothing (see Algorithm 1), and smoothed values were normalized to TPM (UMI-filtered transcripts per million). For visualization as a heatmap in Figure S9a-b, the z-score of every gene across cells was calculated. For scatter plots in Figure S9c-e, the expression of each gene was log_2_(TPM + 1).

### Analysis of scRNA-Seq data using principal component analysis (PCA) and hierarchical clustering

Both PCA and hierarchical clustering were performed on median-normalized and Freeman-Tukey transformed (FT-transformed) data. The procedure that we refer to as “median-normalization” is equivalent to “Model I” in Grün, Kester, and Oudenaarden (2014). It involves first calculating the median total UMI count across all cells in the dataset, and then scaling the expression profile of each cell so that its total UMI count equals this median value. More formally, for a dataset containing *p* genes and *n* cells, let ***u***_*j*_ = (*u*_1*j*_, …, *u*_*pj*_)^*T*^ represent the expression profile (gene UMI counts) of the *j*’th cell (either unsmoothed, or after kNN-smoothing without dividing by k+1). Let *t*_*j*_ = Σ_*i*_ *u*_*ij*_ represent the total UMI count of the *j*’th cell. Then let *t*^med^ = median {*t*_1_, …, *t*_*n*_} be the median total UMI count. Median-normalization then consists of calculating scaled expression profiles 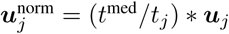.

The Freeman-Tukey transform is a variance-stabilization transformation for Poisson-distributed data proposed by Freeman and Tukey (1950). It is defined as 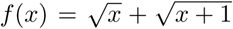. We apply this transformation to the normalized UMI counts to ensure that independently of gene expression level, the absolute level of technical noise is comparable between genes. Specifically, we calculate the transformed UMI counts as 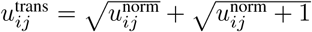.

PCA was performed on median-normalized and FT-transformed data, retaining all genes in the `PANCREAS` and `PMBC` datasets, respectively, using the `sklearn.decomposition`. `PCA` class from *scikit-learn* v0.19.1. Hierarchical clustering was also performed on median-normalized and FT-transformed data, but after filtering for the 1,000 most variable genes, using the `scipy.cluster.hierarchy.linkage` function from *scipy* v1.0.0. More specifically, we calculated the variance for each gene in median-normalized and FT-transformed data, and retained the 1,000 genes with the largest variance. For clustering cells, we used Euclidean distance, and for clustering genes, we used correlation distance. In both cases, we used average linkage. To visualize the clustered data as a heatmap, we re-ordered the genes and cells according to the results of the hierarchical clustering, and standardized the expression values of each gene by substracting the mean and dividing by its sample standard deviation.

### Selection of cell type-specific marker genes

For cell types in the `PANCREAS` dataset, we selected the same genes used by Baron et al. (2016). For the `PMBC` dataset, we manually selected genes based on well-known markers, a previously published analysis of scRNA-Seq PBMC data (Zheng et al. 2017), and literature searches. In particular, for moncoytes, we followed known protein surface markers and selected *CD33*, a myeloid lineage marker, *CD14*, specifically expressed in monocytes, and *CD16*, expressed on a subset of monocytes, as well as certain NK cells and T cells (Naeim et al. 2013). To mark dendritic cells, we selected FCER1A and CLEC9A, both previously shown to be specifically expressed in those cells (Villani et al. 2017). For T cells, we used *CD3D* and *CD3E*, the protein products of which form a dimer of the T cell receptor complex, and are pan T cell markers (Naeim et al. 2013). We also included *CD8A* and *CD8B*, encoding two isoforms of the CD8 T cell co-receptor present on cytotoxic T cells. For NK cells, we included *NCAM1* (CD56), *NCR1* (CD335), and *KLRD1*(CD94), all of which are expressed on NK cells at the protein level (Naeim et al. 2013). Finally, for B cell,s we included *CD19*, *MS4A1* (CD20), and *CD79A*, all well-known B cell markers (Naeim et al. 2013).

### Simulation of scRNA-Seq data

The `SIM-PANCREAS` dataset was simulated based on the `PANCREAS` dataset using the following approach: First, we used smoothing and hierarchical clustering to group the cells in the `PANCREAS` dataset into ten clusters. To do so, we applied kNN-smomothing with *k* = 31. Then, the smoothed dataset was median-normalized, and the normalized values were Freeman-Tukey transformed. Then, the dataset was filtered for the top 2,000 most variable genes, and hierarchical (agglomerative) clustering was performed on the cells, using average linkage and the Euclidean distance metric. The resulting tree was cut at the appropriate height to produce ten clusters. We chose hierarchical clustering over other clustering methods because it simplifies the visualization of clustering results, and because it can ensure a certain degree of consistency between simulated datasets that only differ in terms of the number of clusters simulated.

After assigning all cells to one of ten clusters, we calculated the cluster expression profiles by averaging the expression profiles of all cells assigned to that cluster, using the original (unsmoothed) UMI counts. For each cell in `PANCREAS`, we then simulated a corresponding expression profile for inclusion in the `SIM-PANCREAS` dataset, by looking up the cluster it was assigned to, scaling the cluster expression profile to match the observed number of transcripts for that cell, and then drawing the expression value for each gene from a Poisson distribution with the corresponding *λ* parameter.

To formalize this procedure, let *p* be the number of genes in the `PANCREAS` dataset, and let ***u***_*j*_ = (*u*_1*j*_, …, *u*_*pj*_)^*T*^ represent the expression profile (gene UMI counts) of the *j*’th cell (before smoothing). Let *z*_*j*_ ∈ {1, …, 10} represent the cluster assignment of the *j*’th cell (obtained using hierarchical clustering, as described above). For the simulation, we then define a corresponding set of 10 clusters. Let ***e***_*c*_ = (*e*_1*c*_, …, *e*_*pc*_)^*T*^ represent the true expression profile of the *j*’th cluster, which we define using 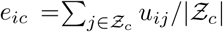. Let *t*_*j*_ = Σ_*i*_ *u*_*ij*_ represent the total UMI count of the *j*’th cell. Let 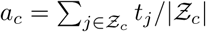 represent the average total UMI count for cells in the *c*’th cluster. We use this information to simulate a dataset with *n* cells. Let 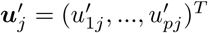, represent the expression profile (gene UMI counts) of the *j*’th cell in the simulated dataset. We obtain each 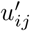 by sampling from a Poisson distribution with mean parameter *λ*_*ij*_, where *λ*_*ij*_ = (*t*_*j*_/*a*_*z*_*j*__) * *e*_*iz*__*j*_.

The `SIM-PBMC` dataset was simulated based on the `PMBC` dataset using a completely analogous procedure.

### Comparison of the accuracies of kNN-smoothing, MAGIC, and scImpute on simulated data

We downloaded MAGIC (commit 4d5efb4) from GitHub, and installed the Python package included. We also installed the scImpute R package (v0.0.4; commit dda0441) from GitHub, using the command `install github`(‱`Vivianstats/scImpute`′). We then applied both methods, as well as kNN-smoothing, to the `SIM-PANCREAS` dataset (testing different parameter choices; see below). For each cell in the dataset, we looked up the identity of the cluster that was used as the basis for the simulation of that cell’s expression profile. The expression profile of that cluster represented the ground truth that the smoothed expression profile should ideally be identical to. To quantify the similarity between the smoothed and the ground truth expression profile, we first applied a log_2_-transformation to both profiles, adding a pseudocount of 1: *f* (*x*) = log_2_(*x* + 1). We then calculated the Pearson correlation coefficient (PCC) between the smoothed and ground truth expression profiles, as well as the root mean squared distance (RMSE) between those profiles. We visualized the results using boxplots in which each value represents the PCC or RMSE of a single profile (cell) after smoothing. We also calculated PCC and RMSE for values transformed using a square root transformation instead of a log-transformation: 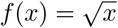, and visualized the results as a boxplot. Finally, we repeated the entire procedure for the `SIM-PBMC` dataset.

For MAGIC, we varied the *t* parameter between 1 and 9, while setting the other parameters to the values recommended in the tutorial provided by the authors of this method: `n_pca_components`=20, k=30, `ka=10`. We reasoned that of all parameters, *t* has by far the strongest effect on the smoothing results, as it is the power to which the Markov affinity matrix is raised. *t* can also be interpreted as the length of a random walk, and larger values of *t* therefore lead to much stronger smoothing (Dijk et al. 2017). For scImpute, we decided to vary both *t* and *K*. In this paper, we refer to *t* as *d*, in order to avoid confusion with MAGIC’s *t* parameter. *d* is the dropout probability threshold that determines the set of genes which will have their expression values imputed. *K* is the number of clusters that determines the sets of candidate neighbors, used to build statistical models to estimate dropout probabilities for each gene (W. V. Li and J. J. Li 2017).

We applied MAGIC using its Python interface (function `SCData.run_magic`), in accordance with the tutorial. We noticed that MAGIC dropped all genes that had no expression in any cell in the simulated datasets, and therefore took care to add these genes back (with zero values) to the smoothed matrix, in order to ensure an unbiased comparison with the other methods (additional or missing zero values change the value of the PCC). We applied scImpute using its R interface (function `scimpute`). It is noteworthy that while the runtime of MAGIC was comparable to kNN-smoothing (usually finishing within seconds or minutes), scImpute routinely took several hours to finish, even when using 4 CPU cores (`ncores=4`).

### Download and combination of 10x Genomics scRNA-Seq data for T cell subsets

We downloaded the following datasets from the 10x Genomics website: “CD4+ Helper T Cells”, “CD4+/CD25+ Regulatory T Cells”, “CD4+/CD45RA+/CD25-Naive T Cells”, “CD4+/CD45RO+ Memory T Cells”, “CD8+ Cytotoxic T Cells” and “CD8+/CD45RA+ Naive Cytotoxic T Cells”. All datasets were first reported by Zheng et al. (2017) and processed by 10x Genomics using the CellRanger software version 1.1.0. For each dataset, we downloaded the “Gene / cell matrix (filtered)”, containing UMI-filtered transcript counts per gene and cell. For each dataset, we then removed genes that were not expressed, removed duplicated genes, and randomly selected a subset of 1,000 cells. We then combined all six datasets into one large dataset containing the union of all the genes from each dataset and all cells. The combined dataset contained 19,208 genes and 6,000 cells.

### Analysis of differential expression between experimentally isolated T cell subsets to identify marker genes

This analysis was performed on scRNA-Seq data form T cell subsets, bead-enriched from peripheral blood (Zheng et al. 2017), and downloaded from the 10x Genomics website (see above). For each T cell subset analyzed (naive CD4 T cells, naive CD8 T cells, and memory CD4 T cells), we sampled a random set of 1,000 profiles from the corresponding dataset, and discarded genes encoding ribosomal proteins, as well as genes located on the mitochondrial genome. To determine a p-value for differential expression for each gene, we applied the Freeman-Tukey transform to all expression measurements, thus stabilizing the technical variance of all expression measurements, and then performed a standard two-sample t-test (assuming equal variance) for each gene. To determine a fold-change value for each gene, we used the untransformed data, averaged the UMI counts for each gene across all cells in each dataset, set all averaged values below 0.05 to 0.05, and calculated log2-transformed ratios for each gene.

### Correlation analysis between smoothed pan-T cell profiles and expression profiles from experimentally isolated T cell subsets

As an external validation of the clustering results obtained for smoothed pan T-cell data, we compared each smoothed expression profile to expression profiles from experimentally isolated T cell subsets. We used the same data by Zhang et al. (2017) as described above (1,000 randomly selected profiles per T cell subset). To generate the subset-specific expression profiles from the scRNA-Seq data, we removed duplicate gene entries, and genes without expression in any of the cells. We then summed up the UMI counts for each gene across all cells, and combined all profiles into a single matrix. For distinguishing naive from memory T cells, this was a matrix containing 17,910 genes and three columns (the profiles for naive CD4 T cells, naive CD8 T cells, and memory CD4 T cells). For distinguishing CD8 and CD4 naive T cells, this was a matrix containing 16,801 genes and two columns (the profiles for naive CD4 and naive CD8 T cells). For each analysis, we then determined the set of genes shared between the matrix with the subset-specific expression profiles and the heatmap containing only the most variable genes from the smoothed pan T-cell data. This was a set of 468 genes for the naive vs. memory T cell analysis, and 191 genes for the CD4 vs. CD8 T cell analysis. We then subsetted the matrix with the subset-specific expression profiles and the smoothed pan-T cell expression matrix using these genes, and transformed the expression values for each genes to z-scores (for both matrices separately). For each subset-specific expression profile, we then determined the Pearson correlation coefficient with all cells in the smoothed pan-T cell data. The results are shown as a heatmap at the bottom of Figure 9a and Figure 9b.

### Measuring the runtime of the kNN-smoothing Python implementations

To measure the runtime of our kNN-smoothing and kNN-smoothing 2 Python implementations, we downloaded the UMI-filtered gene expression matrix of the dataset titled “8k PBMCs from a Healthy Donor” from the 10x Genomics website. After filtering for genes with expression and removing duplicated genes (analogous to our processing of the `PMBC` dataset), we obtained a dataset containing 21,425 genes and 8,381 cells. To test the runtime of kNN-smoothing we randomly sampled *n*=2,000, *n*=4,000 and *n*=8,000 cells (without replacement) and measured the runtime (wall time) of the algorithm for different settings of *k*. For each combination of *n* and *k*, we repeated this procedure three times, and determined average runtimes and standard deviations. The tests for kNN-smoothing were performed using Python v3.5.4, and the tests for kNN-smoothing 2 were performed using Python v3.6.4, both on Ubuntu® 17.10.

## ACKNOWLEDGMENTS

We would like to thank Bo Xia, Maayan Baron, Dr. Gustavo Franc¸a for helpful discussions.

## REFERENCES

Baron, Maayan et al. (2016). “A Single-Cell Transcriptomic Map of the Human and Mouse Pancreas Reveals Inter- and Intra-cell Population Structure”. In: Cell Systems 3.4, 346–360.e4. DOI: 10.1016/j.cels.2016.08.011.

Brink, Susanne C. van den et al. (2017). “Single-cell sequencing reveals dissociation-induced gene expression in tissue subpopulations”. In: Nature Methods 14.10, pp. 935–936. DOI: 10.1038/nmeth.4437.

Cao, Junyue et al. (2017). “Comprehensive single-cell transcriptional profiling of a multicellular organism”. In: Science (New York, N.Y.) 357.6352, pp. 661–667. DOI: 10.1126/science.aam8940.

Carrette, Florent and Charles D. Surh (2012). “IL-7 signaling and CD127 receptor regulation in the control of T cell homeostasis”. In: Seminars in Immunology 24.3, pp. 209–217. DOI: 10.1016/j.smim.2012.04.010.

Dijk, David van et al. (2017). “MAGIC: A diffusion-based imputation method reveals gene-gene interactions in single-cell RNA-sequencing data”. In: bioRxiv. DOI: 10.1101/111591.

Eden, Eran et al. (2009). “GOrilla: a tool for discovery and visualization of enriched GO terms in ranked gene lists”. In: BMC Bioinformatics 10, p. 48. DOI: 10.1186/1471-2105-10-48.

Eisen, M. B. et al. (1998). “Cluster analysis and display of genome-wide expression patterns”. In: Proceedings of the National Academy of Sciences of the United States of America 95.25, pp. 14863–14868.

Fan, Jean et al. (2016). “Characterizing transcriptional heterogeneity through pathway and gene set overdispersion analysis”. In: Nature Methods. DOI: 10.1038/nmeth.3734.

Freeman, Murray F. and John W. Tukey. (1950). “Transformations Related to the Angular and the Square Root”. In: The Annals of Mathematical Statistics 21.4, pp. 607–611. DOI: 10.1214/aoms/1177729756.

Gierahn, Todd M. et al. (2017). “Seq-Well: portable, low-cost RNA sequencing of single cells at high throughput”. In: Nature Methods 14.4, pp. 395–398. DOI: 10.1038/nmeth.4179.

Grün, Dominic, Lennart Kester, and Alexander van Oudenaarden (2014). “Validation of noise models for single-cell transcriptomics”. In: Nature Methods 11.6, pp. 637–640. DOI: 10.1038/nmeth.2930.

Halko, Nathan, Per-Gunnar Martinsson, and Joel A. Tropp (2009). “Finding structure with randomness: Probabilistic algorithms for constructing approximate matrix decompositions”. In: arXiv:0909.4061 [math]. arXiv: 0909.4061.

Hashimshony, Tamar, Naftalie Senderovich, et al. (2016). “CEL-Seq2: sensitive highly-multiplexed single-cell RNA-Seq”. In: Genome Biology 17, p. 77. DOI: 10.1186/s13059-016-0938-8.

Hashimshony, Tamar, Florian Wagner, et al. (2012). “CEL-Seq: single-cell RNA-Seq by multiplexed linear amplification”. In: Cell Reports 2.3, pp. 666–673. DOI: 10.1016/j.celrep.2012.08.003.

Islam, Saiful, Una Kjällquist, et al. (2011). “Characterization of the single-cell transcriptional landscape by highly multiplex RNA-seq”. In: Genome Research 21.7, pp. 1160–1167. DOI: 10.1101/gr.110882.110.

Islam, Saiful, Amit Zeisel, et al. (2014). “Quantitative single-cell RNA-seq with unique molecular identifiers”. In: Nature Methods 11.2, pp. 163–166. DOI: 10.1038/nmeth.2772.

Kharchenko, Peter V., Lev Silberstein, and David T. Scadden (2014). “Bayesian approach to single-cell differential expression analysis”. In: Nature Methods 11.7, pp. 740–742. DOI: 10.1038/nmeth.2967.

Klein, Allon M. et al. (2015). “Droplet barcoding for single-cell transcriptomics applied to embryonic stem cells”. In: Cell 161.5, pp. 1187–1201. DOI: 10.1016/j.cell.2015.04.044.

Kleiveland, Charlotte R. (2015). “Peripheral Blood Mononuclear Cells”. In: The Impact of Food Bioactives on Health. Springer, Cham, pp. 161–167. DOI: 10.1007/978-3-319-16104-415.

Kumari, Sudha et al. (2014). “T cell antigen receptor activation and actin cytoskeleton remodeling”. In: Biochimica Et Biophysica Acta 1838.2, pp. 546–556. DOI: 10.1016/j.bbamem.2013.05.004.

La Manno, Gioele et al. (2017). “RNA velocity in single cells”. In: bioRxiv. DOI: 10.1101/206052.

Li, Wei Vivian and Jingyi Jessica Li (2017). “scImpute: Accurate And Robust Imputation For Single Cell RNA-Seq Data”. In: bioRxiv, p. 141598. DOI: 10.1101/141598.

Love, Michael I., Wolfgang Huber, and Simon Anders (2014). “Moderated estimation of fold change and dispersion for RNA-seq data with DESeq2”. In: Genome Biology 15.12, p. 550. DOI: 10.1186/s13059-014-0550-8.

Lun, Aaron T. L., Karsten Bach, and John C. Marioni (2016). “Pooling across cells to normalize single-cell RNA sequencing data with many zero counts”. In: Genome Biology 17, p. 75. DOI: 10.1186/s13059-016-0947-7.

Macosko, Evan Z. et al. (2015). “Highly Parallel Genome-wide Expression Profiling of Individual Cells Using Nanoliter Droplets”. In: Cell 161.5, pp. 1202–1214. DOI: 10.1016/j.cell.2015.05. 002.

Naeim, Faramarz et al. (2013). “Principles of Immunophenotyping”. In: Atlas of Hematopathology. Academic Press, pp. 25–46. DOI: 10.1016/B978-0-12-385183-3.00002-4.

National Cancer Institute (2017). Division of Cancer Prevention. Accessed March 4, 2002, \url{2018-01-20}. URL: https://prevention.cancer.gov/news-and-events/news/pre-cancer-atlas-pca-and (visited on 01/20/2018).

Paul, Franziska et al. (2015). “Transcriptional Heterogeneity and Lineage Commitment in Myeloid Progenitors”. In: Cell 163.7, pp. 1663–1677. DOI: 10.1016/j.cell.2015.11.013.

Pierson, Emma and Christopher Yau (2015). “ZIFA: Dimensionality reduction for zero-inflated single-cell gene expression analysis”. In: Genome Biology 16, p. 241. DOI: 10.1186/s13059-015-0805-z.

Regev, Aviv et al. (2017). “Science Forum: The Human Cell Atlas”. In: eLife 6, e27041. DOI: 10.7554/eLife.27041.

Risso, Davide et al. (2017). “ZINB-WaVE: A general and flexible method for signal extraction from single-cell RNA-seq data”. In: bioRxiv. DOI: 10.1101/125112.

Rosenberg, Alexander B et al. (2017). “Scaling single cell transcriptomics through split pool barcoding”. In: bioRxiv. DOI: 10.1101/105163.

Sasagawa, Yohei et al. (2017). “Quartz-Seq2: a high-throughput single-cell RNA-sequencing method that effectively uses limited sequence reads”. In: bioRxiv. DOI: 10.1101/159384.

Shekhar, Karthik et al. (2016). “Comprehensive Classification of Retinal Bipolar Neurons by Single-Cell Transcriptomics”. In: Cell 166.5, 1308–1323.e30. DOI: 10.1016/j.cell.2016.07.054.

Subramanian, Aravind et al. (2005). “Gene set enrichment analysis: a knowledge-based approach for interpreting genome-wide expression profiles”. In: Proceedings of the National Academy of Sciences of the United States of America 102.43, pp. 15545–15550. DOI: 10.1073/pnas.0506580102.

Svensson, Valentine, Roser Vento-Tormo, and Sarah A. Teichmann (2018). “Exponential scaling of single-cell RNA-seq in the past decade”. In: Nature Protocols 13.4, pp. 599–604. DOI: 10.1038/nprot.2017.149.

Tang, Fuchou et al. (2009). “mRNA-Seq whole-transcriptome analysis of a single cell”. In: Nature Methods 6.5, pp. 377–382. DOI: 10.1038/nmeth.1315.

The Chan Zuckerberg Initiative (2018). Human Cell Atlas. URL: https://chanzuckerberg.com/human-cell-atlas (visited on 01/21/2018).

Villani, Alexandra-Chloé et al. (2017). “Single-cell RNA-seq reveals new types of human blood dendritic cells, monocytes, and progenitors”. In: Science (New York, N.Y.) 356.6335. DOI: 10.1126/science.aah4573.

Zemmour, David et al. (2018). “Single-cell gene expression reveals a landscape of regulatory T cell phenotypes shaped by the TCR”. In: Nature Immunology 19.3, pp. 291–301. DOI: 10.1038/s41590-018-0051-0.

Zheng, Grace X. Y. et al. (2017). “Massively parallel digital transcriptional profiling of single cells”. In: Nature Communications 8, p. 14049. DOI: 10.1038/ncomms14049.

Ziegenhain, Christoph et al. (2017). “Comparative Analysis of Single-Cell RNA Sequencing Methods”. In: Molecular Cell 65.4, 631–643.e4. DOI: 10.1016/j.molcel.2017.01.023.

